# DNA-RNA hybrid (R-loop): from a unified picture of the mammalian telomere to the genome-wide

**DOI:** 10.1101/2020.08.24.264424

**Authors:** Minoo Rassoulzadegan, Ali Sharifi-Zarchi, Leila Kianmehr

**Affiliations:** Université de Nice, INSERM-CNRS, France; Royan Institute, Tehrān, Iran

## Abstract

Local three-stranded DNA-RNA hybrid regions of the genomes (R-loops) have been detected either by binding of a monoclonal antibody (DRIP assay) or by enzymatic recognition by RNaseH. Such a structure has been postulated for the mouse and human telomeres, clearly suggested by the identification of the complementary RNA TERRA. It was, however, evidenced by DRIP exclusively in human cancer and not in normal human nor in any mouse cell. Based on the observation that in several fractionation procedures, DNA-RNA hybrids copurify with double-stranded DNA, we developed a preparative approach that allows detection of stable DNA-RNA triplexes in a complex genome, their physical isolation and their RNA nucleotide sequence. We then define in the normal mouse and human genomes the notion of terminal R-loop complexes including TERRA molecules synthesized from local promoters of every chromosome. In addition to the telomeric structures since, the finding of the RNA peak, applied in addition to the whole murine sperm genomes, has highlighted a reproducible profile of the R-loop complexes from entire genome suggesting a defined profile for a future generation.

## Introduction

In the recent period, increased attention has been drawn to R-loop complexes, which appear as frequent features of the eukaryotic genomes. These three-stranded structures are made of the two strands of a DNA molecule, one of them displaced by hybridization with a complementary RNA. In addition to short-lived intermediates generated during transcription, detrimental effects of stable local R-loops were documented in several instances, while, on the other hand, they have been considered as potentially important players in genome biology^1-2^. The multiplicity of possible functions is, however, in contrast with the limited number of analytical methods, essentially based on the recognition of the DNA-RNA hybrid either by the monoclonal antibody S9.6, or a catalytic deficient RNase H, coupled with DNA-seq and less frequently, with RNA-seq analysis (reviewed in refs^2-7^). Methodological problems are illustrated by the variety of conclusions reached in different studies^2^.

One physiological instance of such a three-stranded hybrid structure has been proposed for the mouse and human telomeres^6-9^. Following early reports of a positive DRIP assay^10,11^, a consensus seemed established of an R-loop structure involving the terminal DNA repeats of the chromosomes and the complementary TERRA RNA. Evidence was, however, limited to the telomerase-negative human tumor cells (ALT tumors) without any experimental evidence in a healthy human cell, nor in any mouse cell^9^. We developed an antibody-independent assay for detection of stable R-loops. Based on the isolation and sequencing of complementary RNAs copurified with the DNA backbone it can be applied to any biological cell or tissue. First tested on the human and mouse telomeres, it established that in both species, TERRA molecules transcribed from every adjacent subtelomeric promoters are engaged in terminal R-loops. How TERRA example might reflect a stable R-loop structure in general is currently unknown. Here, we study similar research applied to mouse genomes at multiple sites reveals that DRNAs are tightly linked to DNA in complexes sensitive to RNaseH. We integrate the search of enriched peaks from RNA-seq data to discover reproducible DNA/RNA regions throughout the genome. These results also highlight the pairing region of the two sex chromosome (X and Y) which are also present as a DNA-RNA hybrid.

## Results

### Antibody-independent profiling of R-loop complexes: a general approach

We devised a simplified, quick and inexpensive method for detection of RNA-DNA hybrid regions in a complex genome (Supplementary Fig.-1). Unlike the current DRIP assay, it is not based on antigenic recognition of the hybrid, with the advantages of avoiding possible artifact at this level. A variant method was developed starting from the observation that in the most generally used TRIzol chloroform extraction procedure^3,4^, a small but reproducible amount of RNA is retained, together with DNA, at the chloroform-water interface. This material as a whole was dissolved in Tris-HCl 10mM, EDTA 1 mM and ethanol precipitated. The pellet was dissolved in proteinase K hydrolysis buffer and DNA and DNA-RNA complexes were recovered by chromatography on a DNA binding “Zymo-SpinTM” column (Zymo-Research Corp Irvine CA, USA). As illustrated below in the case of the telomeric complex, a specified genomic area could be separately analyzed by the proper DNA cleavage procedure. Alternatively, whole preparations of the DNA-associated RNA were obtained by extensive pancreatic DNase hydrolysis.

### The telomeric TERRA complex

The ends of the human, mouse and yeast linear DNA molecules constitutive of individual chromosomes are long TTAGGG/CCCTAA repeats. After discovery of the homologous UUAGGG-repeated TERRA RNA^5-6^, the DNA telomeric repeats were thought to be engaged in three-stranded R-loops with TERRA. Attempts in normal tissues to directly evidence the R-loop structures have been so far only partially successful. In cell culture mostly from two human pathologies could R-loops be evidence by DRIP assay, namely the telomerase-negative (ALT) cancers^7^ and the ICF syndrome cells^8^. Structure of the telomeres of healthy human cells, as well as of any murine cell, remained to be directly evidenced. One main site of TERRA transcription was identified in mouse and human cells^9,10^ and it has been even considered that transcription from individual subtelomeric promoters on every chromosome is a unique characteristic of a class of malignancies (ALT cancers).

Results were generated in parallel on human and mouse sperm cells. Sperm cells were chosen because the majority of cytoplasmic RNAs is removed at the compacting stage, so that the risk of contamination by cytoplasmic RNAs is minimized. The total nucleic acid fraction recovered from the TRIzol interface after purification was subjected to Msp1 cleavage for resection of the DNA telomeric sequences up to the first upstream CCGG site, less than 2,000 bp from the repeats on every chromosome (see Supplementary Figure 1). After gel electrophoresis in either 8 per cent acrylamide or 1.5 per cent agarose followed by transfer (without denaturation of the nucleic acids) to nitrocellulose for binding of the single-stranded RNA and DNA material, hybridization with oligonucleotide probes complementary to the telomeric repeats revealed a unique and homogeneous signal. Identical results were generated in every cell type tested, exemplified in Figure 1 (Fig.1a-c) for three mouse healthy tissues, brain (B), total testis (T) and purified epididymal sperm (S). In Fig.1a the Telomere probe and Supplementary Table-2 shows the electrophoretic pattern of RNA recovered in the TRIzol aqueous fraction and in Fig.1b, the pattern of the material from the interface DNA-associated fraction (“Tatar-blot” for Telomere associated TERRA RNA). The same gel electrophoretic profiles were generated by probes against the CCCTAA and TTAGGG motifs (Fig. 1b, 1c and Supplementary Table-2). DNA denaturation by NaOH after electrophoresis, a step of the classical Southern blot (see below), is obviously omitted in the Tatar-blot.

**Fig. 1.**
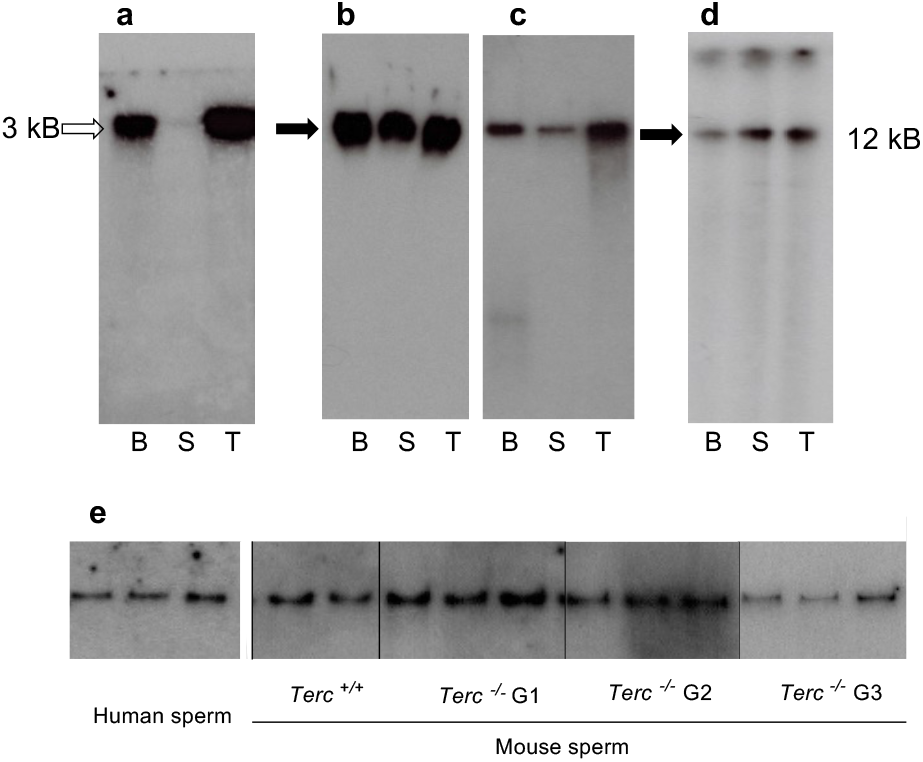
TERRA in sperm and somatic cells. **a**. Northern blot analysis of RNA processed from TRIzol-chloroform aqueous phase, low level in sperm and high level detection of TERRA in Brain and testes, **b and c**. DNA-RNA complexes (Tatar-blot for Telomeres associated TERRA RNA). Whole nucleic acid preparations were analyzed after Msp1-cleavage by electrophoresis on 8% acrylamide gels, nitrocellulose transfer and hybridization with a ^32^P-labelled probe against the (TT/UU)AGGG telomeric motif (b) and complementary sequences (c). **d**. Agarose gel (0,8%) same samples as in b and c. except for a 45 min exposure to 0.5M NaOH and renaturation (standard Southern blot and same probe as in b). In addition to large fragment, a smear of shorter fragments is detected. **e**. Complexes in extracts of human sperm same procedure as in b. Wild type and telomerase-negative (*Terc*^*-/-*^) mouse mutants of the first three generations, which exhibit progressively shortened telomeres. Embryos, testis and sperm biopsies of Terc^−/-^ animals were kindly provided by Drs C. Gunes (Germany) and A. Londono (Paris) and the same results not shown for *Tert*^*-/-*^ animals JAX mice (Jax strain B6, 129S-terttm1Yjc/J, stock# 005423) were purchased from The Jackson laboratory, Bar Harbor.

Specificity was determined by hybridization of the transfers to radioactive probes for other repetitive (SINE, Xist-A) or single copy genomic sequences (Supplementary Table-2), none of them generating a significant signal (data not shown). Electrophoretic migration of the complexes, equivalent to that of a high molecular weight linear DNA molecule (Fig.1 panel b), was clearly not informative as to the actual size. When the same extracts were further processed for Southern blot analysis by denaturation in 0.5 N sodium hydroxide, transfer and hybridization (Fig. 1d) revealed molecules with a smeary profile starting with the expected large size of the telomere (>12 kb).

Complexes with the same electrophoretic profiles were identified in comparable amounts in every mouse cell tested, from embryonic fibroblasts to adult tissues (testis, brain, liver, kidney) as well as cultivated cell lines and short-term cultures of differentiated cells, as exemplified by mouse brain extracts in Fig. 1b. This was also the case of human tissues (Supplementary Table 1), including saliva, blood and sperm (Fig. 1e). Identical electrophoresis profiles were generated for murine and human sperm extracts by probes against the CCCTAA motive (Fig. 1e). These complexes do not appear to depend on the telomerase, as they were observed in three generations of telomerase-negative mouse cells (both *Tert-/-* and *Terc-/-*, Fig. 1e). In the case of the *Tert-/-* and *Terc-/-* mutant, they could also be evidenced in embryo fibroblasts culture from homozygous crosses during the first three generations in which the mutants maintain telomeres^11,12^ (data not shown) and after immortalization in culture of mutant cell lines (MR, unpublished).

Properties of the complexes were compatible with the R-loop structure schematically drawn in Figure 2 (Fig. 2a), the G-rich strand of telomeric DNA displaced by the hybrid of TERRA (Fig. 1b and d). Nucleolytic attack was performed by incubating extracts (20 µl) for 30 to 60 min with either RNaseA 0.5 µg/µl, RNaseH 1 u/µl, or DNase 10 u/µl (Fig. 2b). The electrophoretic signal disappeared after incubation with DNase, pancreatic RNase, formamide, or RNaseH, while it was not affected by RNase A (Fig. 2b). Another feature of the complex was indicated by a slower migration rate in gel electrophoresis performed in the presence of 20µM PhenDC3(4), a reactant of G quadruplexes^13^ (Fig. 2b). Visualization of the complex requires resection from chromosomal sequences. The clearest results were generated by cleavage with Msp1 at sites (CCGG) present at least once in the immediate subtelomeric 1-2Kb of every mouse and human chromosome, so that the contribution of chromosomal sequences is minimal (facilitated transfer on the nitro cellulose membrane). Other restriction endonucleases tested (BamH1, Bgl2, Alu1, Mse1, and Dpn1) did not generate comparable results, most probably because they cleave at greater and variable distances from the telomeres, thus generating larger double stranded fragments that are not retained efficiently on nitrocellulose. In agreement with previous observations of increased methylation of CpG sites in subtelomeric regions^14^, cleavage with Hpa2, a methylation-sensitive isoschizomer of Msp1, failed to evidence the complex (not shown).

**Fig. 2.**
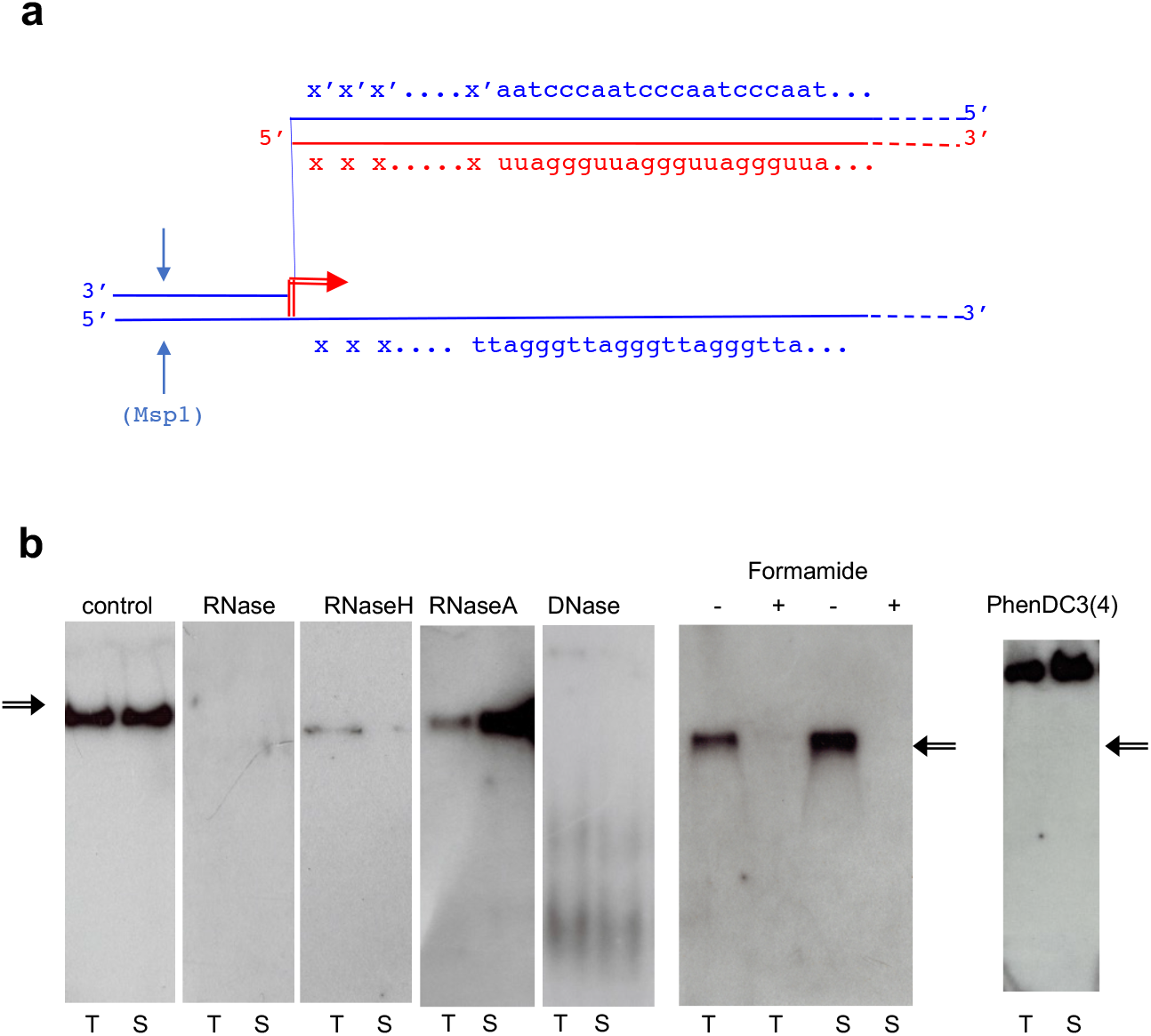
The telomeric TERRA complex. **a**. A simplified R-loop model. TERRA RNA (red line) is assumed to originate from a subtelomeric promoter (red arrow); blue lines: telomere and sub-telomere DNA; “xxx” : 5’ TERRA sequence from subtelomeric DNA (xxx/x’x’x’); arrows: Msp1 cleavage. **b**. Complex detection Tatar blot (Telomeres associated TERRA RNA) performed as in Fig. 1b on the indicated mouse cells (T: testis and S: sperm). Left to right: detection of the (TT/UU)AGGG and CCCTAA motifs (see Fig. 1b). Then complex evidenced in several conditions (control), in extracts treated with pancreatic (DNase free RNase ref. 11119915001) RNase 0,5 µg/µl, RNaseH 1u/µl (ref. M0297S), RNaseA 1u/µl (ref. 101091690 01) and DNase 10u/µl, respectively (all provided by Roche Life Science), and (right) electrophoresis after addition of either 50% formamide (denaturation conditions+/ or -) or 20 µM phenylDC(3)4 (right).

The same results were also generated in parallel on a series of human and mouse cells, *ex vivo* organ extracts most of them normal cells, but including cultured cell lines (Supplementary Table-1), and, as a positive control, the human U-2 OS cancer cell line (ATCC® HTB-96(tm)) in which DNA-RNA telomeric hybrids have been previously identified by DRIP^7^.

### Transcription of individual telomeres from local subtelomeric promoters

A key point was to establish whether the observed structure is present in only one or a few telomeres, or, to the contrary, is a general feature of the genome. It has been generally assumed that R-loops are generated by transcription from adjacent promoters. On the other hand, a major part of TERRA has been reported to be transcribed from chromosome 18 in mouse cells and from chromosome 20q in human^9,10^ so that our data so far are compatible either with a unique R-loop on these chromosomes or with integration in trans in other chromosomes of RNAs from the mouse 18 and human 20q loci. To the contrary, however, it appeared that unique subtelomeric sequences of different chromosomes are represented in the DNA-associated fraction. We generated probes for sequences between the first Msp1 site and the G-rich repeats of the telomeres of chromosomes 9, 17, 18, 19, X and Y telomeres and probes for the complementary strand up to the C-rich repeats (indicated “xxx / x’x’x’” in Fig. 2a). By using UCSC genome browser Blat tool^15^, we ensured that these flanking regions are unique in the whole genome. All the probes (Supplementary Table 2) generated positive hybridization signals in the complex as illustrated in Fig. 3a for the subtelomeric sequences of chromosomes 17, 18 and Y. As controls (not shown), none of the probes for sequences on the centromere side of the terminal Msp1 sites generated any signal.

**Fig. 3.**
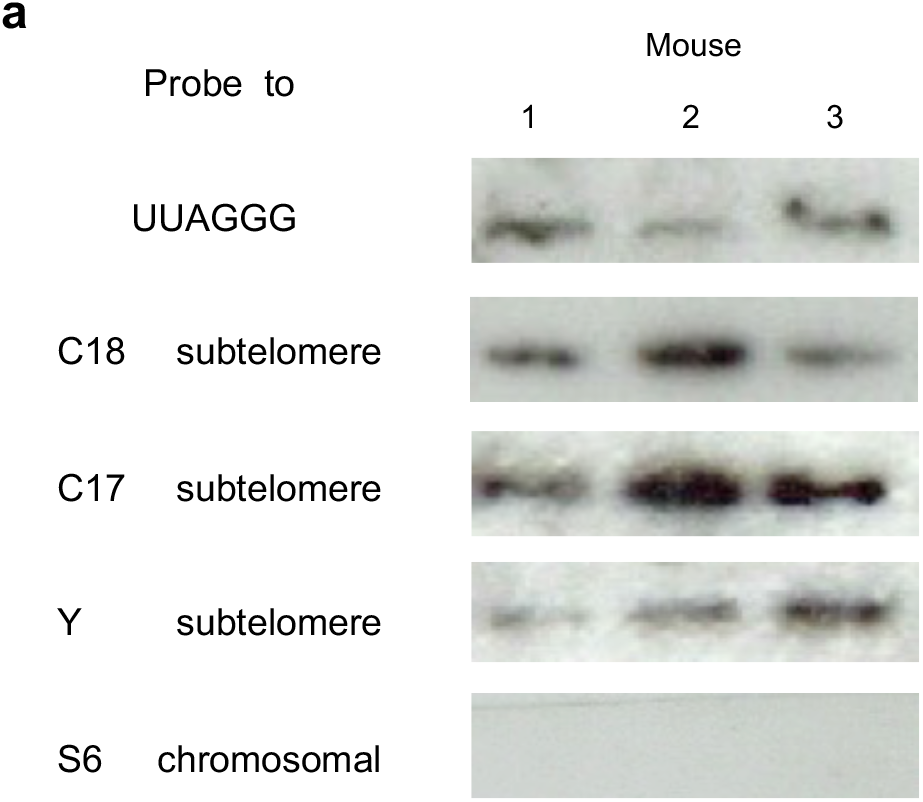
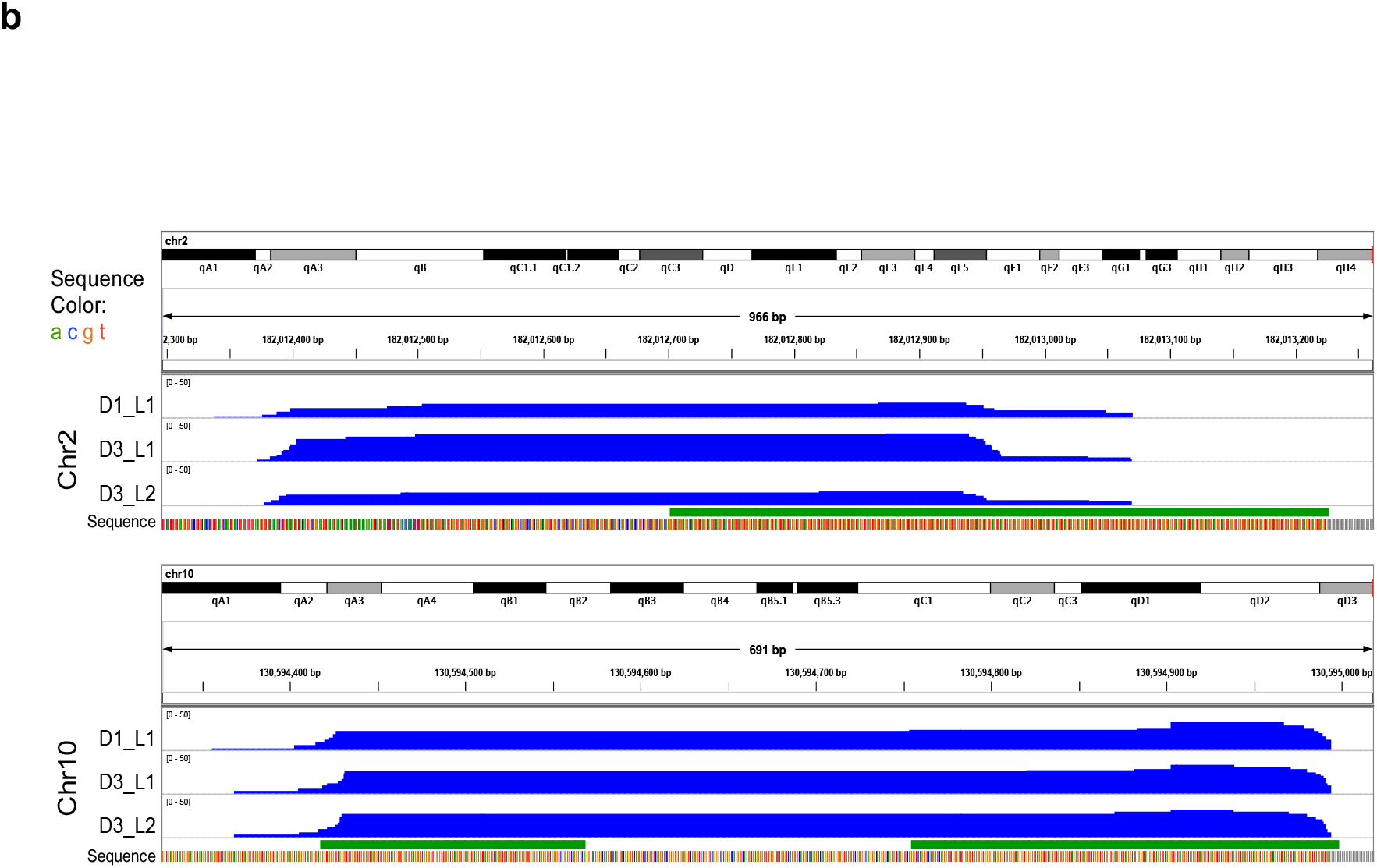
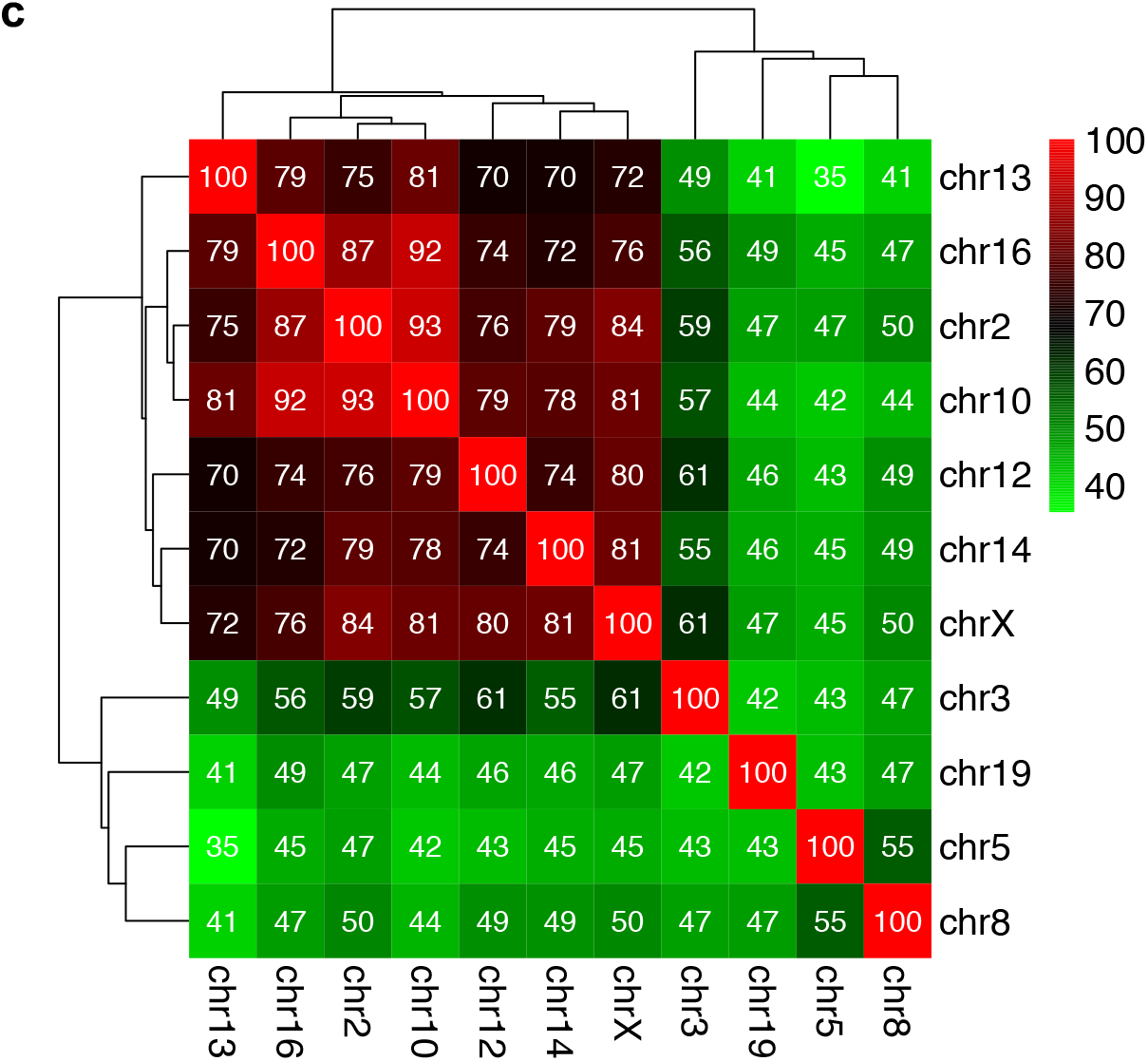
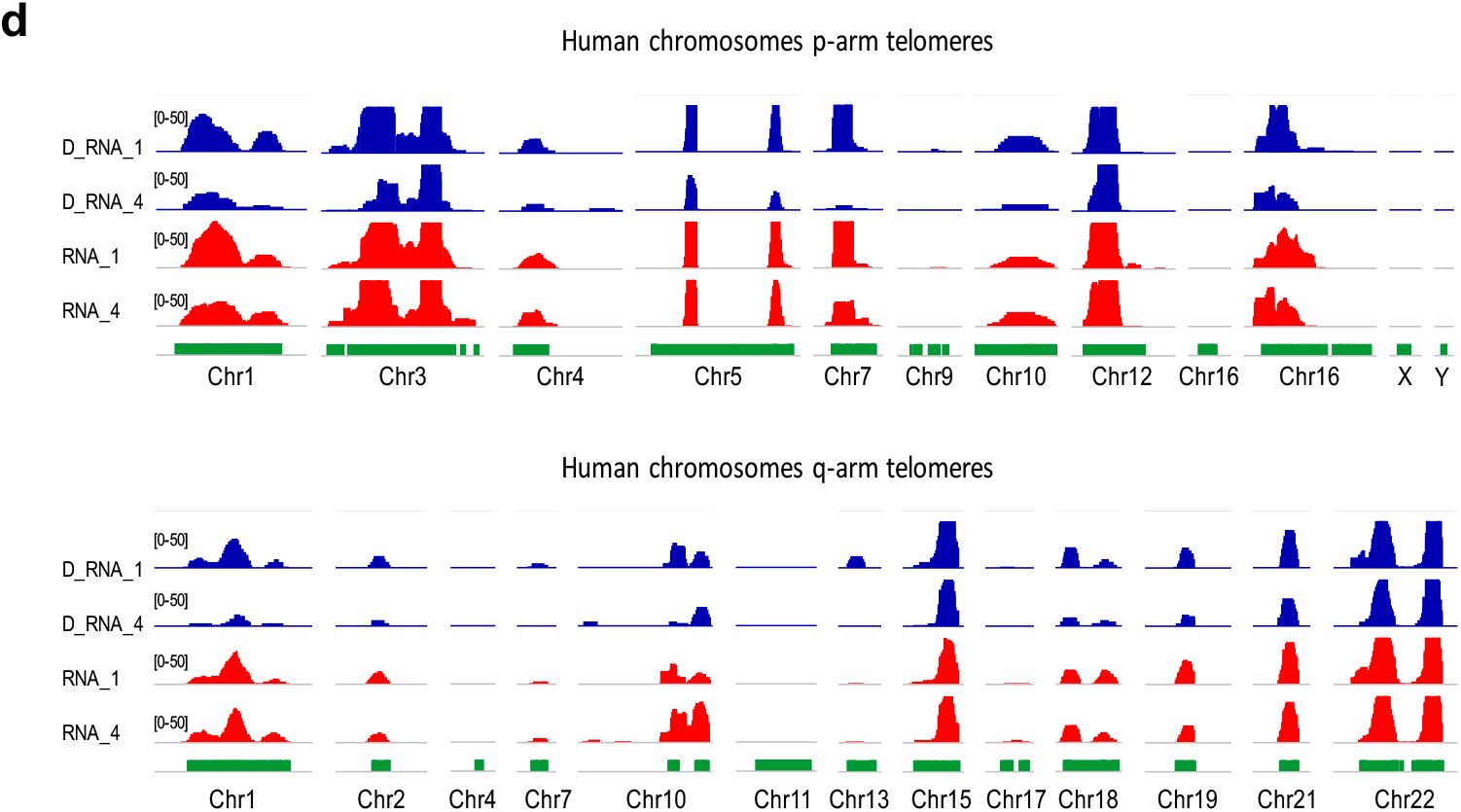
TERRA transcripts originate from subtelomeric promoters. **a**. After electrophoresis and transfer of complex of mouse sperm performed as in Fig. 1b Tatar blot (Telomeres associated TERRA RNA), hybridization with probes for (top to bottom) UUAGGG repeat (8-fold), subtelomeric sequences of chromosomes 18, 17 and Y (schematized as xxx in Fig. 2a), and S6 ribosomal RNA as a negative control. The chromosomal sequences were randomly taken from the mm10 mouse genome assembly and checked in each case to be unique in the whole genome. Left to right: analysis performed on males of three different strains, *C57, Balb/c* and *B6D2*. DNA-bound RNAs were prepared from epididymal sperm and from the total testis (estimated 60 per cent of diploid cells) of a 6 month, a 14 month-old male and two sperm samples from man at reproductive age, all showing identical electrophoretic profiles of the DNA-RNA complexes (Tatar-blot not shown). Mouse spermatozoa were collected from the cauda epididymis and ejaculated human sperm kindly provided by the Fertility Clinics of the University. In both cases, standard protocol is already reported previously and Kianmehr & al. ^24^ summarized: motile spermatozoa were recovered and washed MEM buffer (1 mM Na pyruvate, 0.5 mM EDTA, 50 U/ml penicillin, 50 mg/ml streptomycin, and 0.1% BSA) by centrifugation at 3000 rpm. Sperm pellets were resuspended in phosphate-buffered saline (PBS) and centrifuged again. The pellets were washed twice in 50 mM HEPES buffer pH 7.5, 10 mM NaCl, 5 mM Mg acetate, and 25% glycerol. Samples of human sperm were identically processed. TRIzol extracted interface materials were ethanol precipitated and followed by overnight incubation at 56°C in Tris buffer 20 mM pH8, EDTA 50 mM, with 0.5% SDS, 20 µM dithiothreitol and 400 µg/ml Proteinase K. Total nucleic acids after enzymatic removal of the proteins was followed by fractionation of RNA and DNA-bound RNA. Extracts were fractionated by the ZR-Duet™ DNA/RNA MiniPrep Plus protocol according to the specifications of the manufacturer (www.zymoresearch.com) see Supplementary Fig. 1. In all cases, the fraction that remains bound to the DNA further purified after DNase digestion. About 10-100 ng of RNA were analysed by high throughput sequencing on Illumina HiSeq 2500 or Illumina MiSeq (Eurofins Medigenomix GmbH, Ebersberg, Germany). They (Eurofins) generated libraries designated D1 and D3 from the DRNAs of a 6 month (D1) and a 14 month-old male (D3), with R1 and R3 the corresponding nucleoplasmic sperm RNAs. Totally, 66 millions (m) reads were mapped to unique sites in the mouse genome (D1), 69m (D3), 77m (R1) and 87m (R3). From the total testes it was prepared only the DRNA libraries TD1, (70m reads) and TD3 (82m), and from human sperm of a healthy unknown donor, the hD 1 and 4, 50m (reads) and hR 1 and 4 (88m). Average length of the reads was about 100 nt. All the primary sequence characteristics for sample libraries of the DNA-bound RNA and free-RNA sequences of germ cells are summarized in Supplementary file 2 (Table A). Sperm and testes transcripts profiling and alignment-based visualization were performed for analysis. Comparison of transcript abundance between samples as measured by RNA-seq yielded TPM (transcript per million) values for the different samples. With FastQC version 0.11.5 quality control of the RAW sequencing reads was performed. Trimming of bad-quality reads was performed using Trimmomatic version 0.36^27^. Based on the FastQC results 10 nucleotides were cropped from the 5’ end of each read, and trimmed bases with Phred score lower than 20 from heading and trailing of each read. Also the Illumina adapter sequences were removed, and the trimmed reads with a size less than 30 bp were removed. Hisat2 was used to align trimmed reads to the reference assembly (GRCh38 for human and GRCm38 for mouse)^27^, with quality, alignment and plot-bamstats utilities of samtools^28^. HOMER version 4.9 was used for peak finding and generating bedGraph files for visualization of alignment^29^. The bedGraphs were cross-sample normalized to have 107 reads per sample in order to make visualization of different samples comparable. Visualization of alignment was performed using IGV version 2.3.67^30^. Genome-wide locations of TERRA repeats were found by aligning sequences of length 24 and 48 composed of 4 and 8 copies of TERRA to the reference genome using bowtie2 using “-a” argument to report all alignment^31^. We also detected statistically significant peaks of expression using HOMER (4-fold expression over local region, Poisson p-value over local region < 0.0001, FDR adjusted p-value < 0.001). We retrieved the sequences of those peaks located in telomeric regions using UCSC genome browser. Multiple sequence alignment of these peaks was performed using EMBL-EBI Clustal Omega^35^ and using R package heatmap was visualized percent identity of the alignment on matrix. **b**. Expression patterns of mice DNA-bound RNA samples over q-arm telomeric regions of chromosomes 2 and 10. The upper track of each panel shows the whole chromosome, and the tiny red bar to the right is the focused genomic region. The blue tracks show expression signals in 3 samples (normalized read frequencies per 10^7^ reads). The green track shows location of TERRA sequences in mice genome (minimum four consecutive TTAGGG repeats). The nucleotide sequence is also shown at the bottom of each panel with color bars (the grey bars show undetermined “N” nucleotides). **c**. Heatmap showing the percent identity matrix between each pair of sequences of chromosomes12 and 10. **d**. Sperm RNA-seq signals over TERRA sequences at p- and q-arm telomeric regions of several human chromosomes. Each track shows a different sperm sample (D_RNA_1 & 4 are DNA-bound RNAs, and RNA_1-4 are free cytoplasmic RNA samples). The heights of the peaks show normalized expression level (number of reads per 10^7^ reads) over each genomic region, with equal 0-50 scale. The green track shows location of TERRA telomeric repeats in human genome (minimum four consecutive TTAGGG repeats). All human telomeric regions with a sequenced TERRA sequence in human reference assembly GRCh38 are shown.

We further ascertained by Illumina high-throughput analysis of the DNA-bound RNA molecules that their sequences are compatible with transcription in each telomere from a local promoter Figure 3 (Fig. 3 and Supplementary File 2). Starting from purified mouse sperm, we first confirmed by RNA-seq that a fraction of the DNA-associated RNA molecules amounting to 0.5 to 2 per cent of the reads showed the characteristic UUAGGG TERRA repeats (minimum four repeats, Supplementary Fig.3). We then searched the RNA sequence libraries for chromosomal sequences next to the repeats and extended the search in the 5’ direction to collect upstream flanking regions. We again ensured that these flanking regions are unique in the whole genome. As exemplified for chromosomes 2 and 10 in Fig. 3b, their 5’ extension in the mouse genome indicated that TERRA expression originates in each case from promoters in the immediate subtelomeric region. Complementary evidence for the synthesis of these RNAs from distinct promoters is shown in Fig. 3c with the heatmap showing the percent identity matrix between each pair of sequences of chromosomes 2 and 10. The maximum identity is 93% making thus impossible that these reads could originate from a unique chromosome. The available sequence data and libraries allowed us to reach the same conclusion for chromosomes 2, 3, 5, 10, 12, 13, 14, 16, 19 and X see Supplementary Fig. 2. R-loop structures involving the products of local transcription therefore appear as a general feature – especially taking into account some constraints in the analysis, for instance the fact that in the latest mouse assembly (mm10), chromosomes 4, 6, X and Y remain to be sequenced up to the repetitive telomeric tract. Testes samples TD1 and TD3 also contain TERRA in their DRNA fractions as shown in Supplementary Fig. 2.

### Extension to the telomeres in human sperm: TERRA transcripts in the individual telomeres from local subtelomeric promoters

We then asked whether subtelomeric transcription may occurs also from every chromosome in human. Then, we assessed RNA sequencing from human sperm of healthy unknown donors, see methods for preparation RNAs from attached to DNA and free fractions. Interestingly, DRNA fraction from human sperm as in mouse show hybrids telomeric TERRA RNAs Fig. 3d RNA-seq signals over TERRA sequences at p- and q-arm telomeric regions of several human chromosomes. Each track shows a different sample (hD_RNA_1-4 are DNA-bound RNA samples, and hRNA_1-4 are cytoplasmic RNA samples). The heights of the peaks show normalized expression level (number of reads per 10^7 reads) over each genomic region, with equal 0-50 scale.

TERRA RNAs resulted from every telomeric end in mouse and human normal cells is consistent with idea that TERRA RNAs is expressed from subtelomeric promoters and remained associated to the telomeres ends. Our results has to be confronted with the previous reports of major TERRA promoters in C18 (mouse) and 20q (human) chromosomes^9,10^. There is in fact but an apparent contradiction since these promoters were identified in experiments in which TERRA was prepared by standard TRIzol extraction from abnormal cells such as cancers cells, and thus corresponds to the nucleoplasmic RNA fraction thought to be involved in other, non telomeric functions^16,17^. Thus, it is important to recover TERRA RNAs from both fraction nucleoplasmic and DNA associated.

### Extension analysis to other DRNAs with UUAGGG motif repetition

Since the association of TERRA transcripts with telomeres could be evidenced by the proposed method, we also asked if TERRA hybrids located at defined non-telomeric sites could be detected. In order to try to distinguish stable complexes of “R-loop” ^18,19^ and the transient association of the nascent RNAs the framework of the process of transcription, we carried out a possible verification in the data from DRNA-seq of mice as above (mouse sperm and testes). In Supplementary Figure 3 for a general evaluation of the RNA molecules associated with DNA (indicated “DRNAs”), the RNA-seq analysis has been extended to the entire genomic UUAGGG repetition containing transcripts in mouse sperm, in the total testicular tissues (indicated D1 and D3, and TD1 and TD3 in the Supplementary Figure 3).

Analysis of entire genome heatmap in Figure 4a shows representative landscapes of RNA-seq of chromosomes 2, 10, 12 and 18. For each chromosome, the green strip shows the location of TERRA-homologous regions (minimum 4 consecutive TTAGGG repeats) which are present at several chromosomal locations in DRNA fractions.

**Fig. 4.**
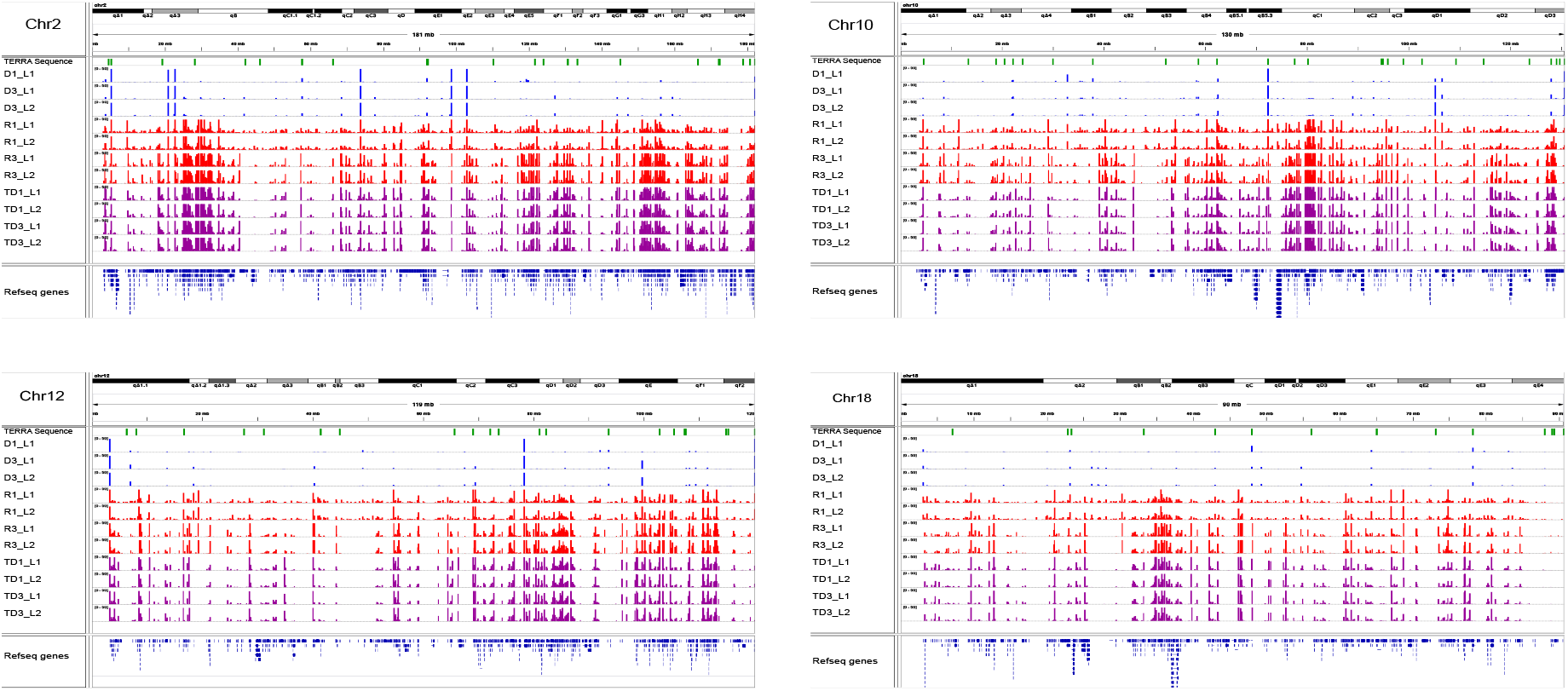
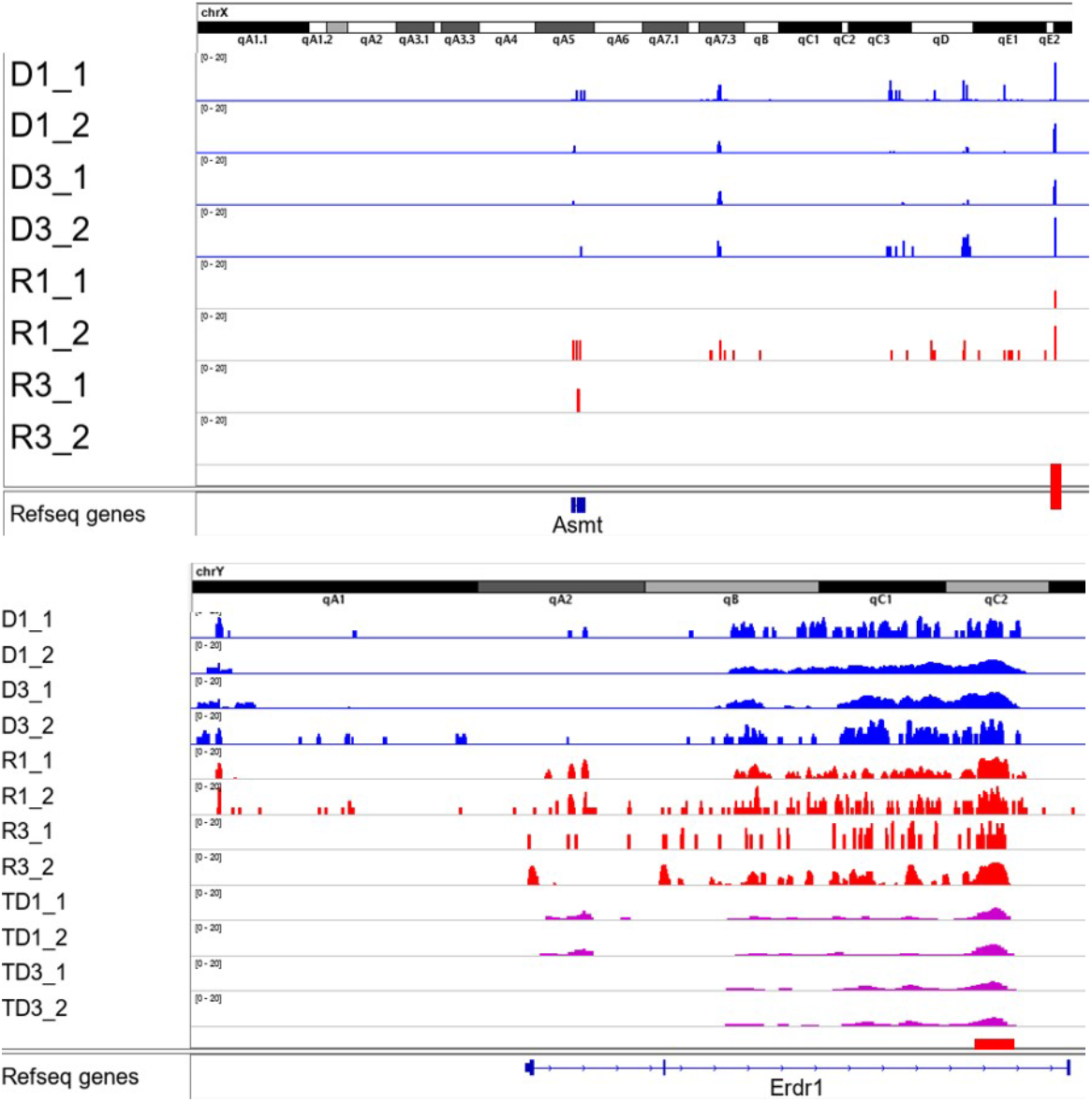
Examples of expression landscapes of the whole mice chromosomes 2, 10, 12 and 18 and from the pseudoautosomal regions of the two sex chromosomes. For each chromosome, the green track shows the location of TERRA-homologous regions (minimum 4 consecutive TTAGGG repeats). Figure 4a, the following tracks show different sperm samples (D1-D3 are DNA-bound RNA, R1-R3 are cytoplasmic RNA, and TD1-TD3 are from testes DNA-bound RNAs). The heights of the peaks show normalized expression level (number of reads per 10^7 reads) over each genomic region, with equal 0-50 scale. Known genes in RefSeq database are located at the bottom of each panel. Figure 4a, TERRA RNA spots are present in sex chromosomes (see Supplementary Figure 4). The number of TERRA RNA foci colocalized with pseudoautosomal regions for instance, Asmt and Erdr1 according to Lee et al. ^17^.

**Fig. 5.**
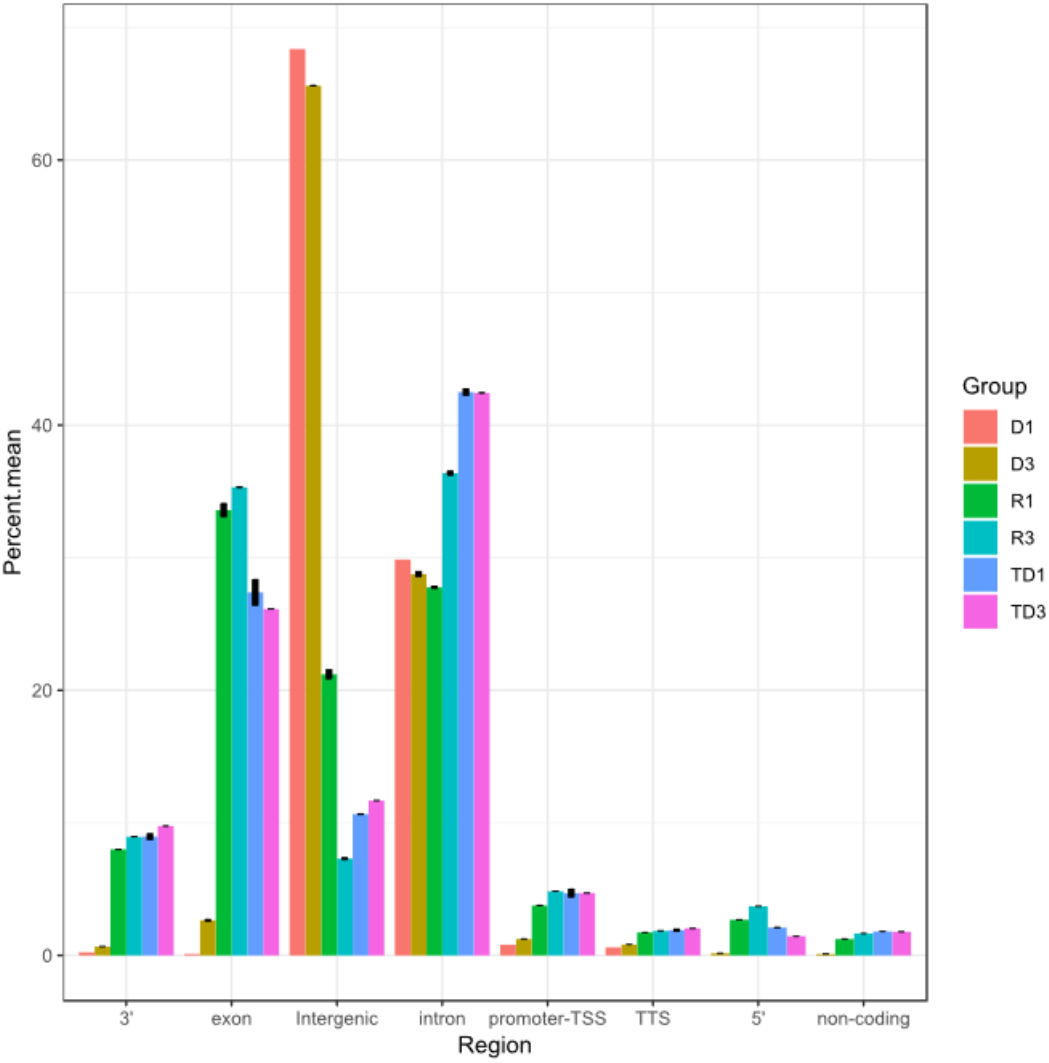
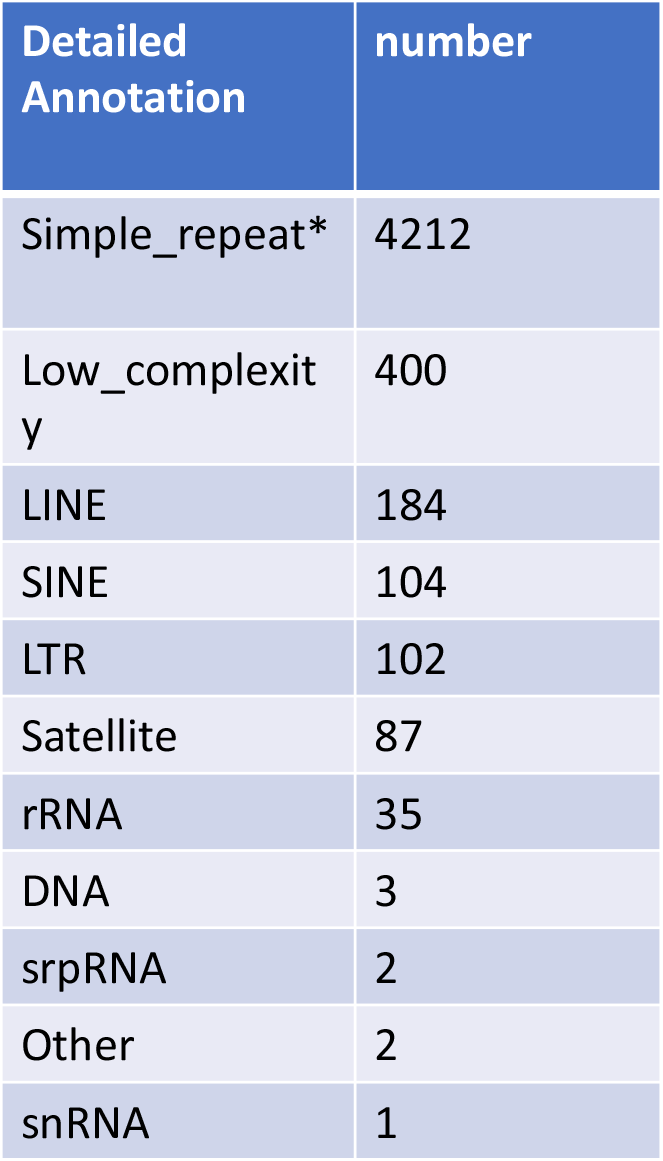
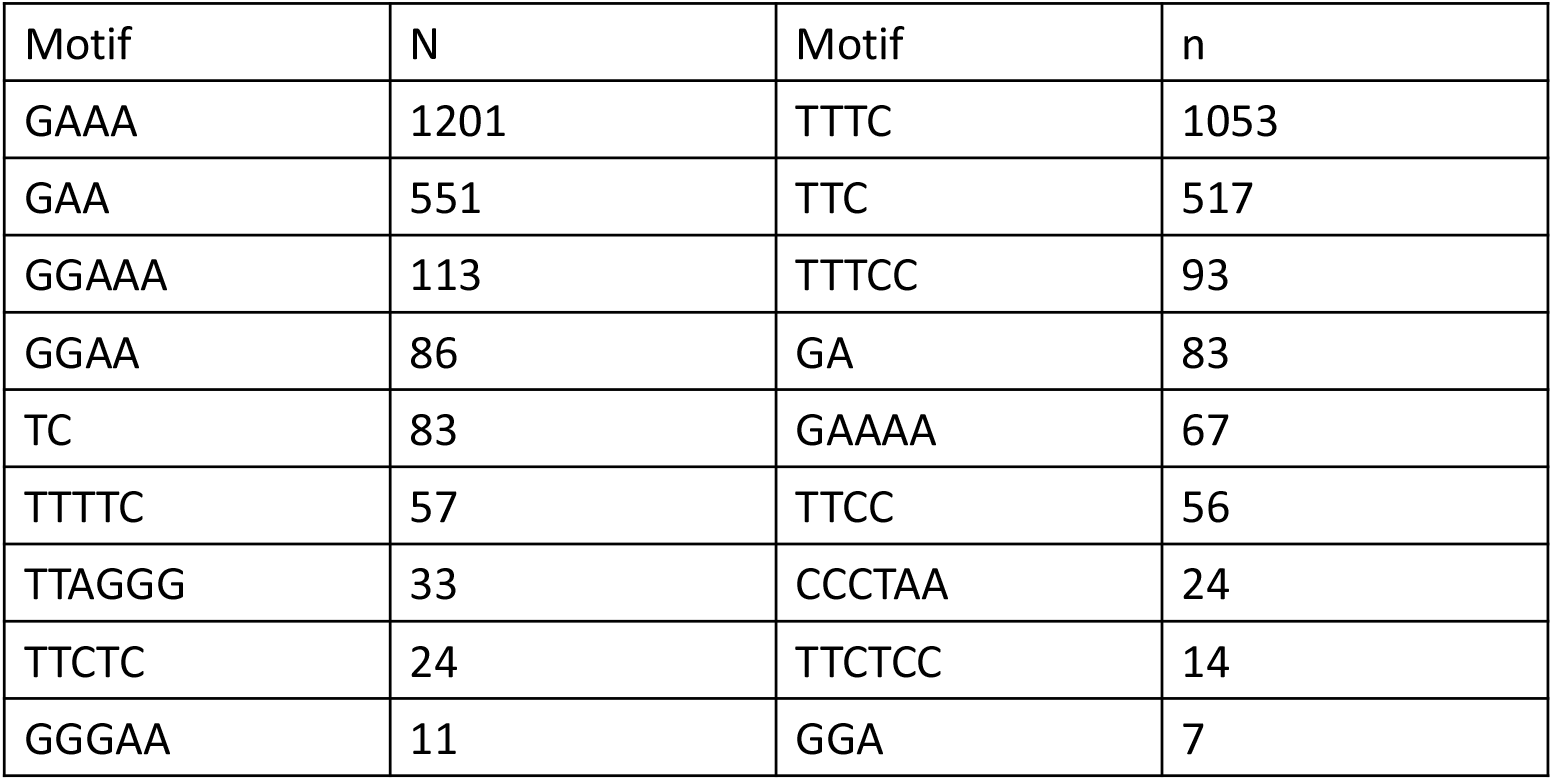
Total transcripts analysis for peak finding on entire genome. **a**. A false discovery rate (FDR) cutoff of 0.001 was used for identifying significant peaks. Peak annotation was performed using Homer annotate_peaks to associate each peak with nearby genes and genomic features. Annotated positions for promoter-TSS, exons, introns and other features was based on the mm10 transcripts. Enrichment analysis of associated genes was also performed using Enrichr for identifying Gene Ontologies (GOs) and pathways significantly over-represented by associated gene sets. For each sample group (D=DNA bound RNA in Sperm, R=free RNA in Sperm, TD=DNA bound RNA in Testis), we measured percent of significant peaks that were located in each genomic region. Y-axis shows average percentage in biological replicates of each sample group. Error-bars represent standard deviations of replicates. **B**. Short list of various motif repeat.

In addition, research on TERRA has identified a fraction of TERRA transcripts near the sex chromosome that undergo pairing in both sex, the inactive X chromosome (Xi) of female cells and the Y ^17^ chromosome (see Supplementary Figure 4 a and b). Exemplified transcripts are visualized (Figure 4b) within the pairing center in the testes and sperm RNAs data. As shown in the Figure 4b Erdr1 locus with UUAGGG repeats was present from the Y chromosome. On the other hand, the Asmt transcripts are present on X chromosome. These results indicate a sample retention of PAR-TERRA transcripts in pseudoautosomal regions of X (Asmt) and Y (Erdr1) in the DRNA fraction of the mouse sperm and testes.

### DRNA is not restricted to the UUAGGG repeats

RNA-seq analysis of sperm and testicles of mice reveals transcripts from the whole genome ref. Here we have analyzed data only from mice because DRNA-seq data is available for two types of samples: sperm (haploid) and testis containing both haploid and diploid cells (germ and somatic cells). A general overview of RNA associated with DNA revealed, in addition to TERRA, a wide variety of sequences, as shown for mouse sperm and testes in Fig. 4. In the sperm DRNA fraction only a few significant peaks are observed at several sites in the genome and are also present in all samples (R and TD) with high reproducibility while a number of low signals are present along of all chromosomes, specific to the types of preparation (homologue between D or homologue between the R fractions). To further characterize these relevant peaks we have performed the search for sequence homology, these sequences are transcribed from different genomic regions see Supplementary Table 3 with an average of 1kb and do not contain sequences or homologous motifs.

To further test the possibilities offered by the analysis, we compared the distribution between the DRNA of the main hybrid regions and the unbound rRNA between sperm and somatic (testis) cells. The annotated positions of the promoter-TSS (transcription start site), TTS (transcription termination site), exons, introns and other characteristics were based on mm10 transcripts. In Figure 6 (Fig.6a) we see a higher level of intergenic transcripts in sperm DRNA samples compared to all others. While sperm-free RNA and testes DRNA are distinguished in the exon regions. In the intergenic region, only a single repetition is observed (Fig.6b). In Supplementary Table 4 the enriched transcripts are listed in different regions. These results clearly reveal that in addition to TERRA, other transcripts are associated in the structure of the R-loop hybrids. Preferential intergenic transcripts retention in sperm DRNA fractions are still unknown but cannot be ignored in future research plan.

## Discussion

Here, we have (i) demonstrated that a fraction of TERRA RNA is stored packed *invivo* with the genome in the form of DNA/RNA hybrids in the telomeres of each chromosome in sperm (mouse and human) and others mouse organs (testes, brain, and liver) (ii) found that non telomeric TERRA RNA repeats at different genomic level are also attached to the DNA (iii) identified that a fraction of the TERRA repeat containing transcripts near the sex chromosome are attached as DRNA to chromosome Y or X and comes from separate genes (iv) define a general profile of transcripts attached to DNA throughout the mouse genome and discover especially important difference in intergenic transcripts retention in sperm DRNA fraction and the different proportion of groups of transcripts attached to DNA between sperm and testicular cells.

We propose that the hybrid DNA/RNA interaction in semen would serve to preserve RNA signalling and could easily be extracted to be identified by RNA-seq analysis and further tested in functional tests. DNA associated RNA (DRNA) extraction is straight forward without need of a supplementary interacting guide molecules (antibody, gene product), cheap, highly reproducible and efficient with bioinformatics analysis to generate genome-wide picture of DNA/RNA hybrids transcripts. Our data shows that all of these stable R loops could provide insight into the amount of RNA maintained in a silent state in the sperm genome. These transcripts are of interest in view of the various reports highlighting a role for sperm RNAs in the epigenetic control of gene expression^20-21^.

The parallel is possible. With S9.6, RNase H and more recently MapR^22^ established that the GC content is important in the recognition of the R-loop by antibodies and in particular by RNaseH^23^. Alternatively, here unlike the previous methods, R-loop detection without protein of DNA/RNA hybrids in normal cells (sperm and testes) reveals, the stable structure of DNA-RNA hybrids which are not particularly rich in CpG. The two types of interaction are not mutually exclusive. However, additional factors must be involved to select or keep the signals. Our study suggests that the difference between antibody-directed (complex) and DNA-RNA hybrids is that, although both appear to be R-loop detection, the latter apparently despite the fact that are highly specific does not involve any other mechanisms, these are direct DNA/RNA interactions.

In the testes, several studies are already reported genome-wide transcripts and it is also assumed that from last stage of elongated spermatids, with elimination of the majority of cytoplasmic RNAs, genome-wide transcripts remain in the sperm. We have recently reported profiles of DRNA-seq mouse sperm transcripts from all coding and non-coding regions of the genome^24^. Contrary to the analysis of the previous individual transcripts, we have here carried out a peak annotation in order to associate the location of peaks/region. This was to assign maps in terms of genomic characteristics.

### Genomic profiles

The purpose of the region in the R-loop is very variable and debated. Our current data provide clear evidence of stable region of the R-loop. In addition, our study shows that sperm (silent cell) in the DRNA fraction reserve transcripts ratios different from the free fraction in sperm and testes DRNA (active cell). Although spermatozoa do not express transcripts they must still, in a manner still unknown select transcripts to be transported. Sperm (DRNAs fraction) have a significantly higher level of intergenic transcripts than exon, intron, 3’, 5’, promoter-TSS (transcription start site), TTS (transcription termination site) and non-coding transcripts. These DNA-RNA hybrids regions detected here consist mainly of short, simple repeat sequences. Others repeated elements such as LIN, SIN LTR and satellite are also present but less frequently. Intergenic transcripts have different characteristics from mRNA and are often involved in chromatin remodelling, transcription control, including enhancer or silencing activities. Therefore, a difference in the level of DNA/RNA hybrid in different regions in particular a higher level in the intergenic region compared to other regions, can be determined by the state of the R-loop of the spermatozoa. Although the accumulation of R-loops is described with genomic instability, here, the DNA-RNA hybrid detected at the physiological level could have a role in cellular characteristics, serving as signal to keep cells in the active or inactive stage. In human spermiogenesis, the large abundance of intergenic RNAs is reported^25^ but now to clarify their roles more functional research is needed. In addition, a recent study suggests that R-loops are involved in determining cell fate ^26^.

### Genomic profiles of the TERRA (telomeric and non-telomeric)

DRNA-seq analysis applied to TERRA complexes of the human and in particular mouse telomeres provides significant new information. The fraction of the telomere associated with DNA synthesized from promoters adjacent to all chromosomes end, is demonstrated in human cells and in a variety of mouse cells and tissues, a conclusion that could not be completely reached with DRIP-seq analysis. Molecular analysis evidences DNA-RNA hybrids at the level of telomeres are maintained in all types of cells. In addition, RNA-seq analysis in the testes, sperm of mice and human sperm confirmed that the DNA-RNA hybrids (telomeres) transcribed in the testes of all chromosomes are maintained until the final stage of maturation in the epididymis. TERRA RNA of the sperm is thus transferred to the offspring with genomic DNA.

The feasibility tested on the analysis of TERRA associated with telomere is also used to identify from the sex chromosomes PAR-TERRA (UUAGGG)n containing transcripts previously established in ES cells^17^ here in mouse DNA-RNA hybrids sperm fraction. The difference in the DRNA transcripts stored between Y and X in the PAR-TRERRA linked to sex is intriguing. One wonders why sperm contain DRNA from PAR-TERRA transcripts. From now on, one objective will be to understand that the DRNA linked to sex chromosome can influence the activities of the X and Y chromosome reaching the oocyte.

Evidence of DNA-RNA hybrids for both TERRA and generalization to genome-associated transcripts in semen underscore ongoing questions about how these hybrids might influence genetic and/epigenetic memory? Among several interpretations, one possibility is a phenomenon unique to sperm: the RNA produced by the spermatid at the time of conversion of the chromatin to its final compact structure can remain associated with the locus, perhaps only the last oligonucleotide segment synthesized, giving the artefactual image of a complete RNA taking into account the sequencing procedure. However, in a way in active testicular cells DNA/RNA hybrids are also observed. In addition, a noticeable difference in the peak ratios of different region between sperm and testes are visible. More than a still preliminary observation against this interpretation is the fact that our less advanced analysis of somatic cells also shows a hybrid DNA/RNA structure detectable with the same method at the telomeres. These studies demonstrate that, RNA appears strongly maintained in a stable complex DNA-RNA hybrid structure and other studies may reveal that its meaning and function may be intentional and selective for a given cell.

In conclusion, we believe that the approach designed to identify the telomeric TERRA has wider applications. These results are clearly encouraging and should be extended to different processes and diseases. Future research is needed to reveal specificities, biogenesis and functions of RNA associated with DNA.

## Acknowledgments

We are grateful to E. Gilson, C. Gunes and A. Londono for the gift of biopsies of telomerase-negative mice and to S. Muller (Paris) for the gift of PhenDC3(4). This work is supported by grant 2019-2020 of La Fondation Nestlé France to Minoo Rassoulzadegan.

## Supplementary material

**Supplementary Table 1.**
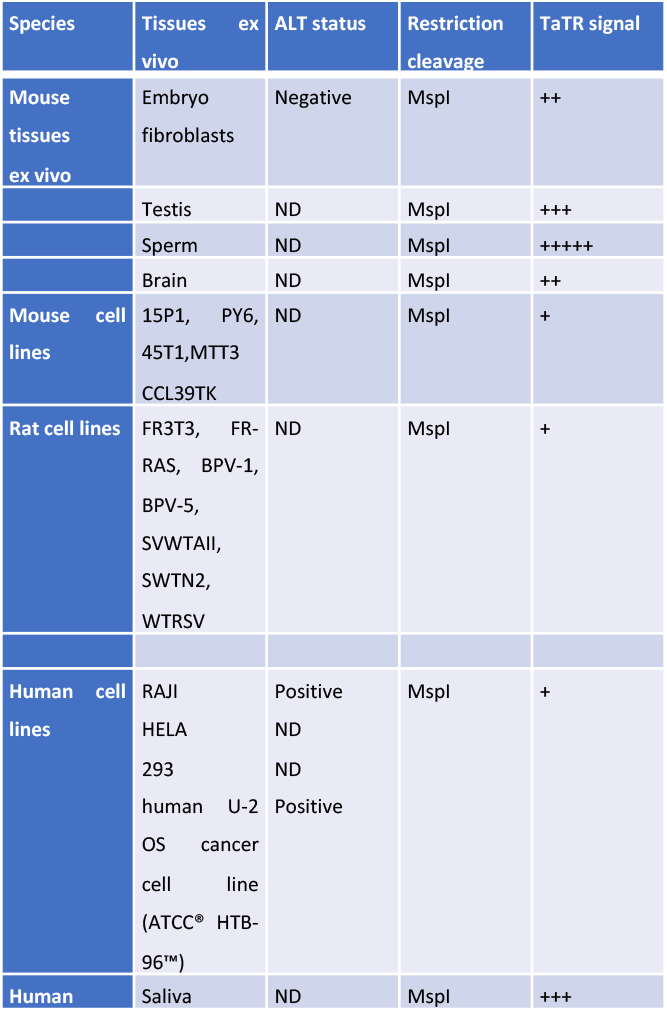
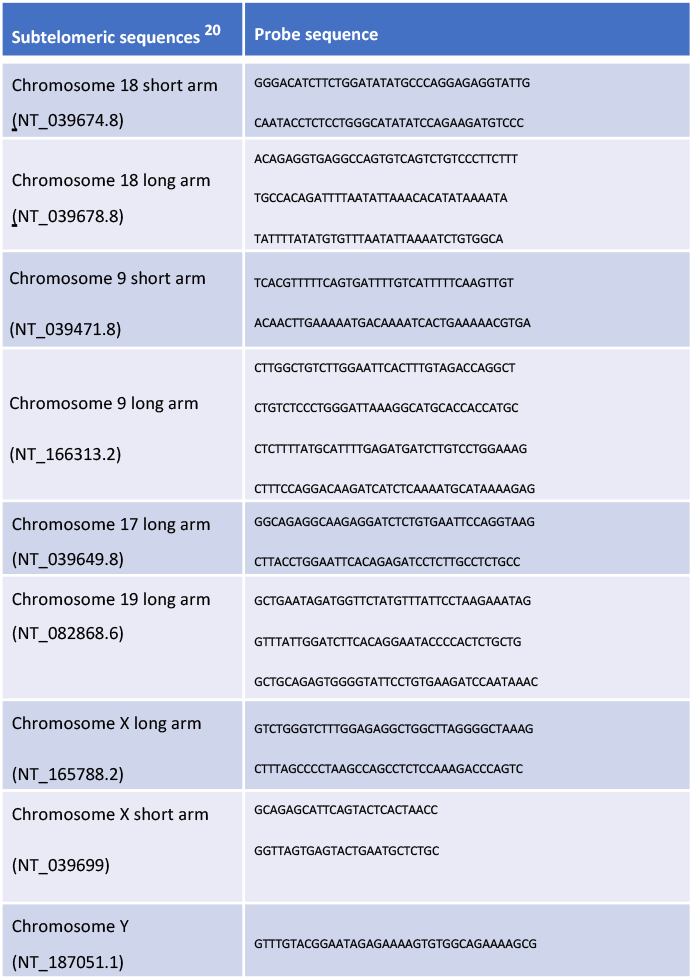
A sample of a total of more than 20 cell lines tested for detection of telomeric TERRA complexes showing different species and telomerase status.

**Supplementary Table 2.**
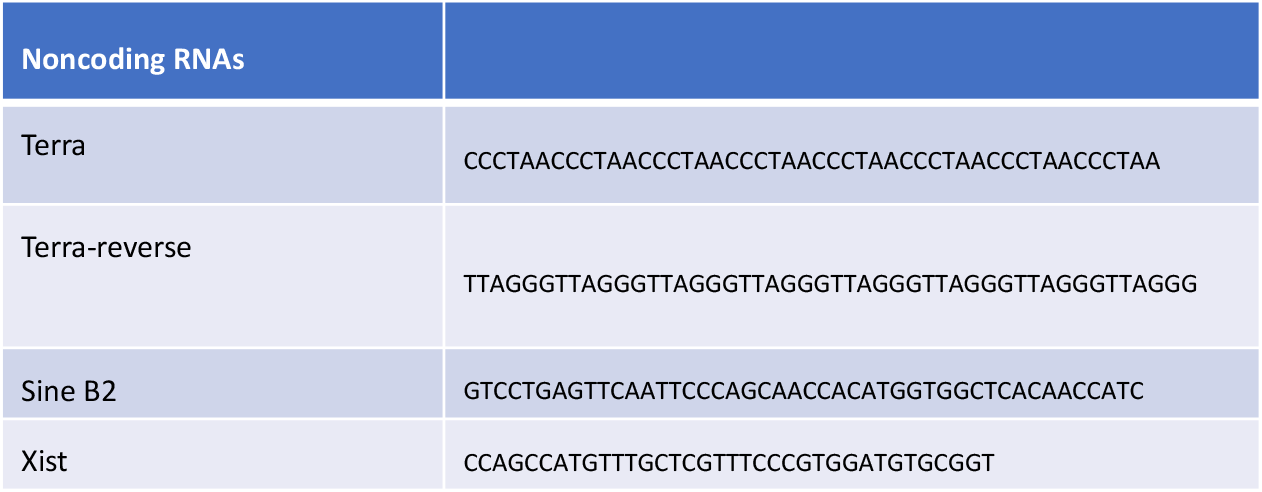
Deoxyribonucleotide probes.

**Supplementary Table 3.**
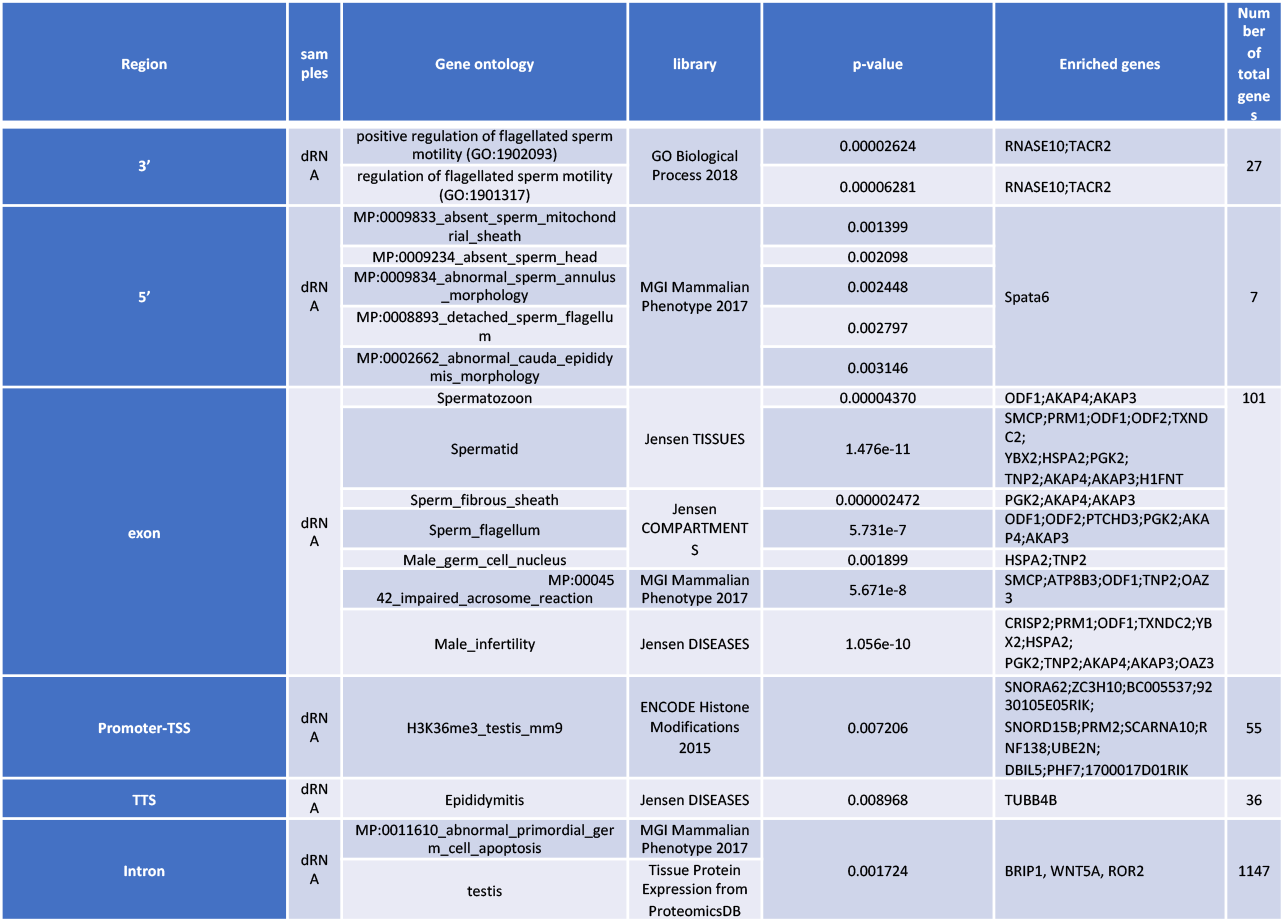
Characterize feature name, length, position and retrieved sequence of different transcripts for a single region of each rare high-peak signals which are presented in the sperm DRNA (blue star peak in Fig. 4) fraction as well in free-RNA fraction and the testes DRNA for sequence homology.

**Supplementary Table 4.**
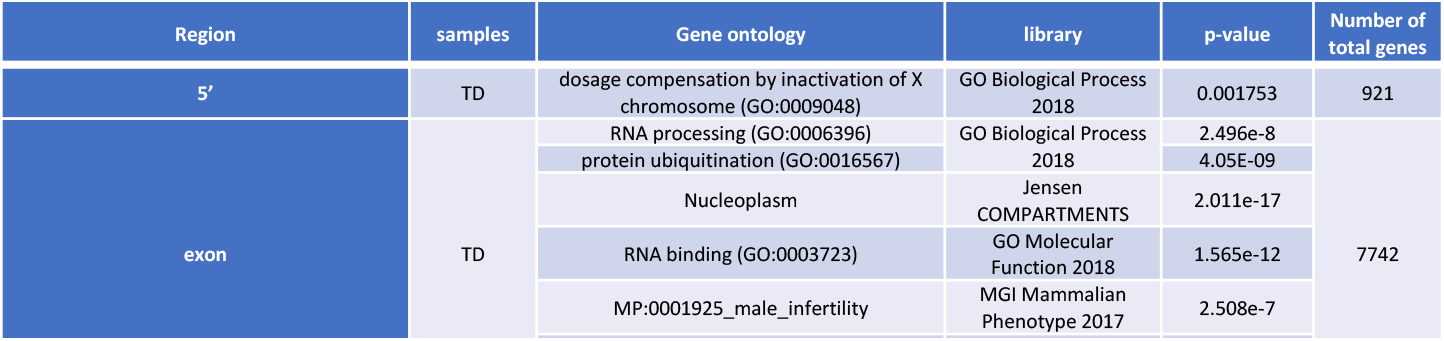
Gene set analysis of different regions using Enrichr tool in DRNA from sperm and testes.

**Supplementary Fig. 1.**
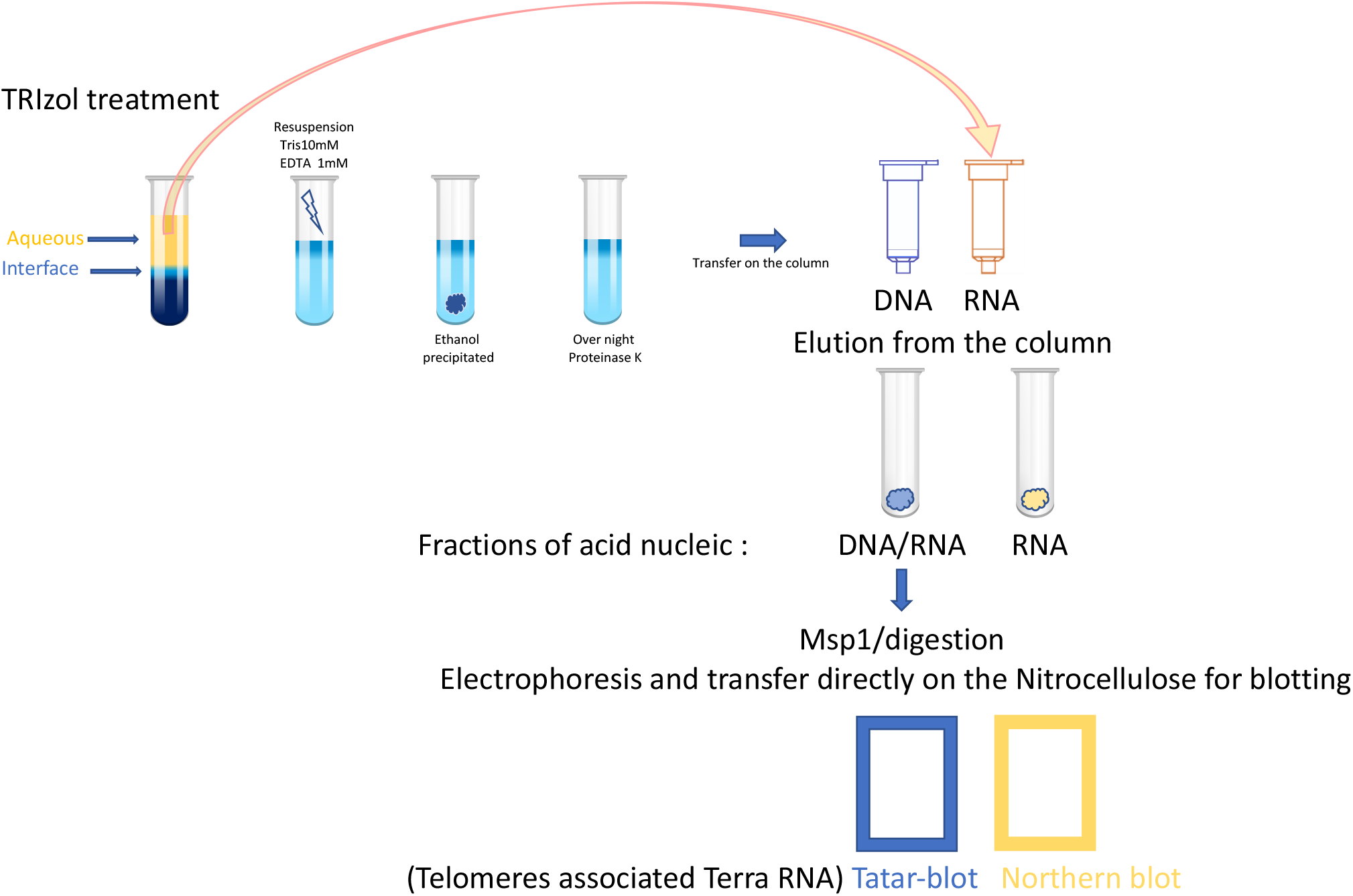
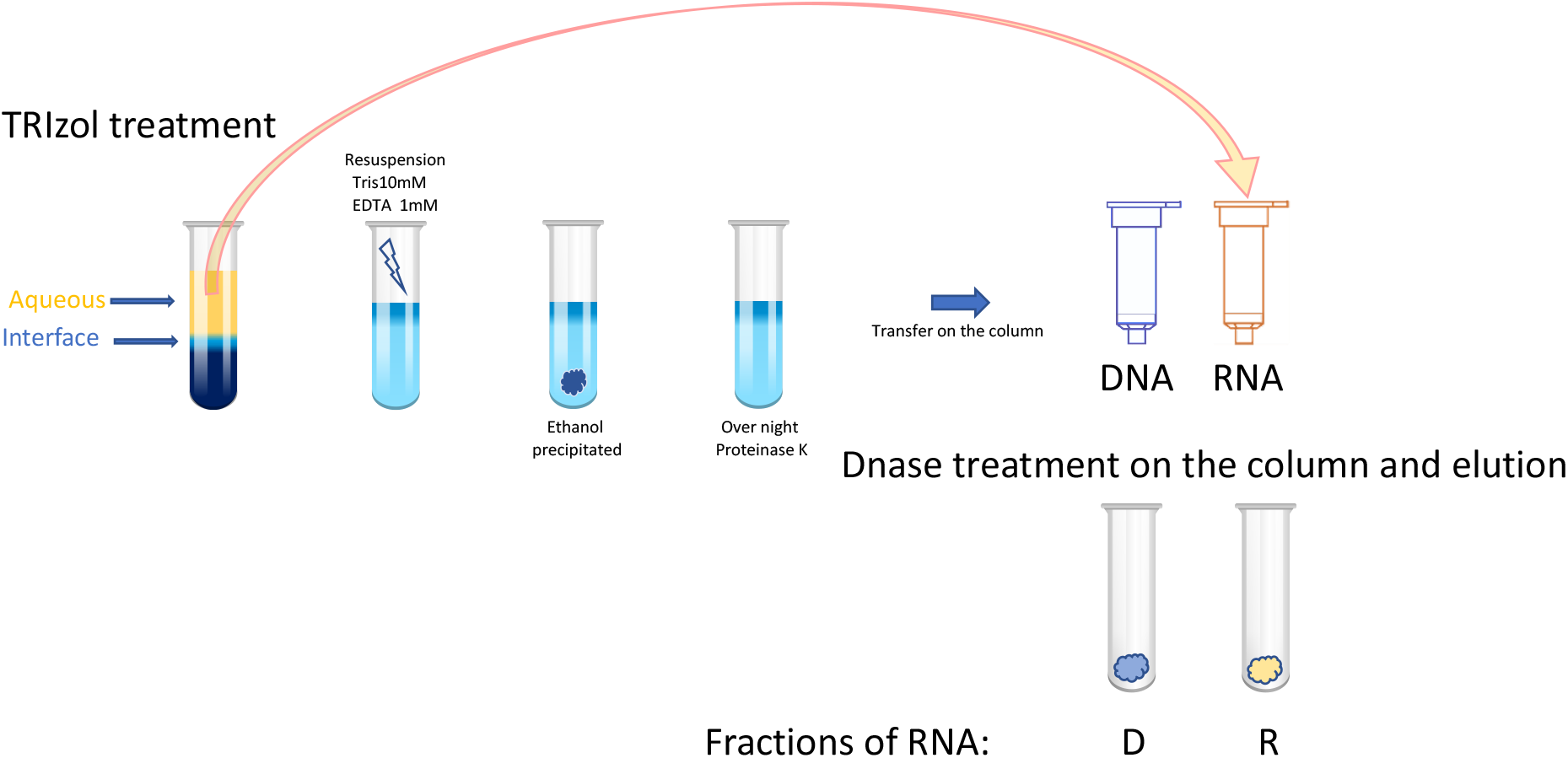
Schema of the nucleic acid extraction protocol.

**Supplementary Fig. 2.**
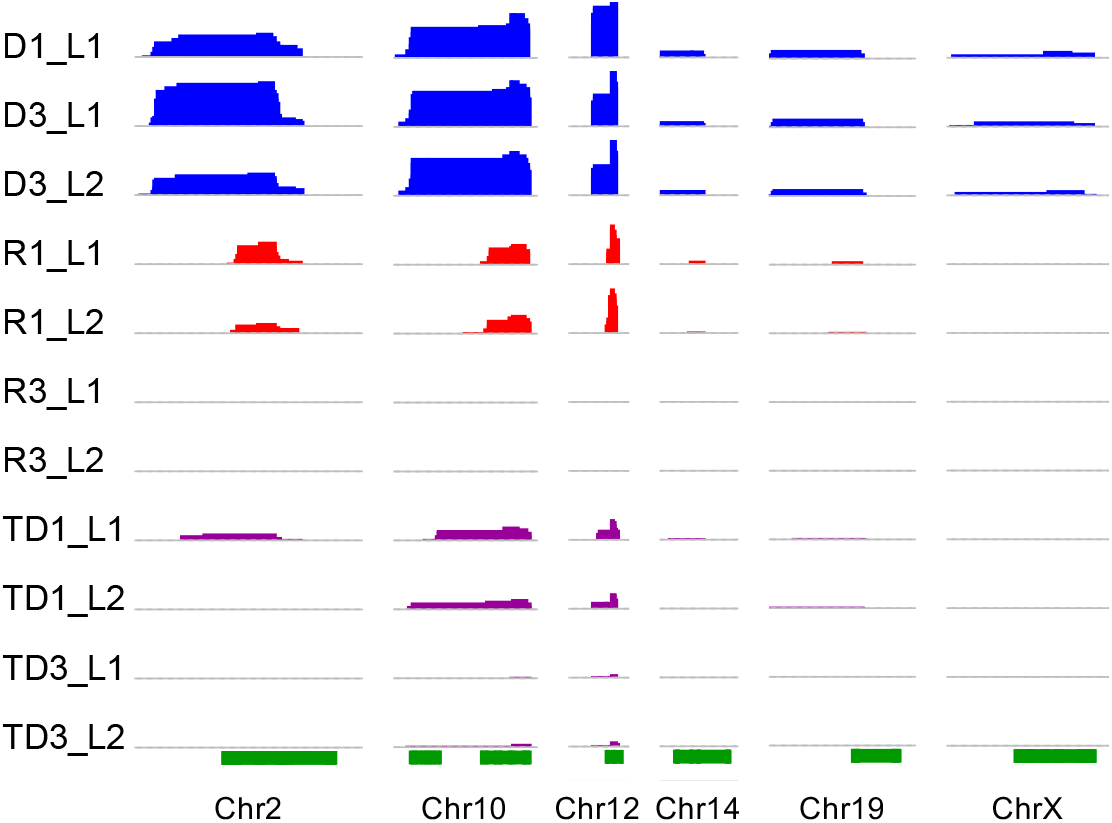
RNA-seq signals over TERRA sequences at q-arm telomeric regions of several mice chromosomes. Each track shows a different sample 1 (male of six months) and 3 (male of 14 month) D1-D3 are DNA-bound RNA, R1-R3 are cytoplasmic RNA from sperm, and TD1-TD3 are DNA-bound RNA from testes. The heights of the peaks show normalized expression level (number of reads per 10^7^ reads) over each genomic region, with equal 0-50 scale. The green track shows location of TERRA sequences in mice genome (minimum four consecutive TTAGGG repeats).

**Supplementary Fig. 3.**
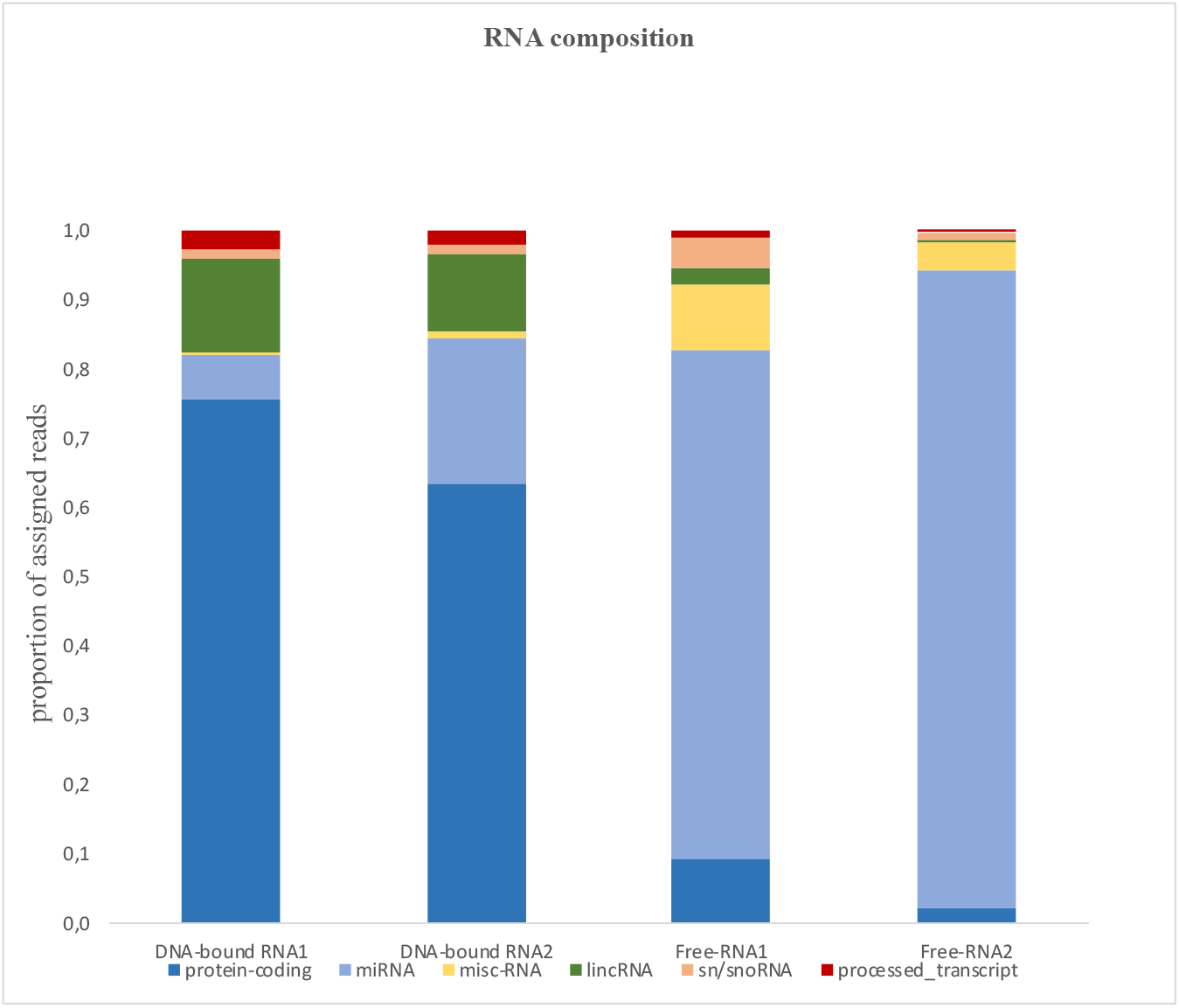
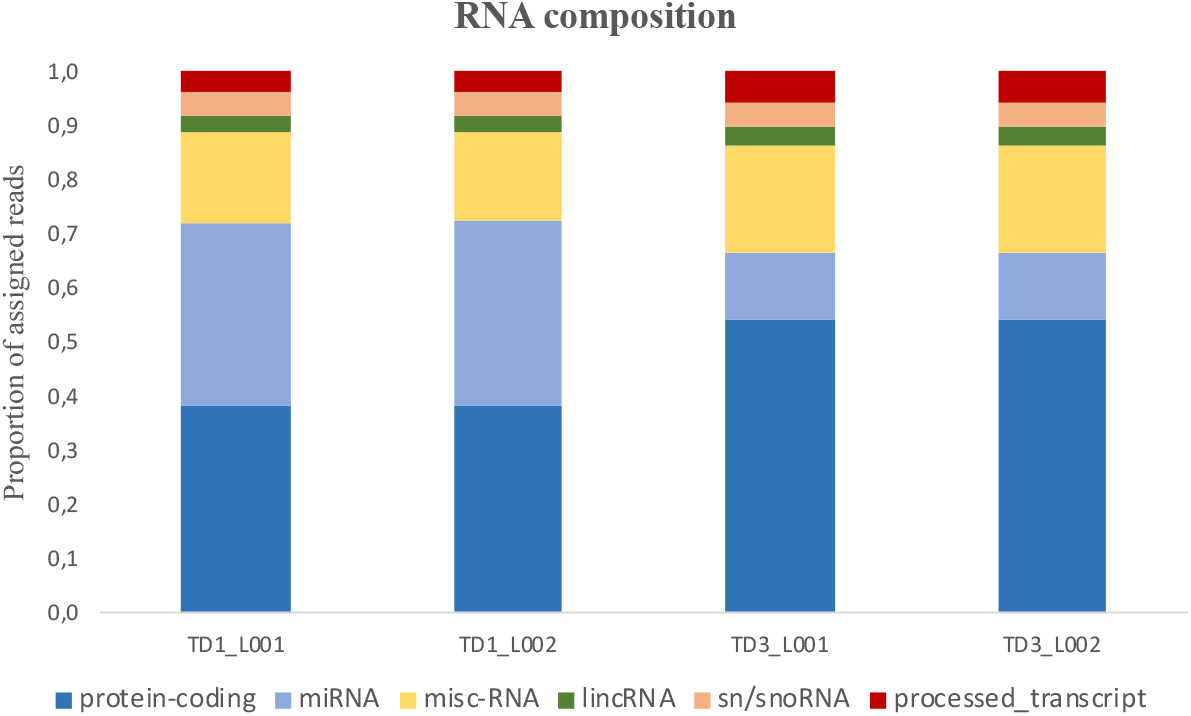
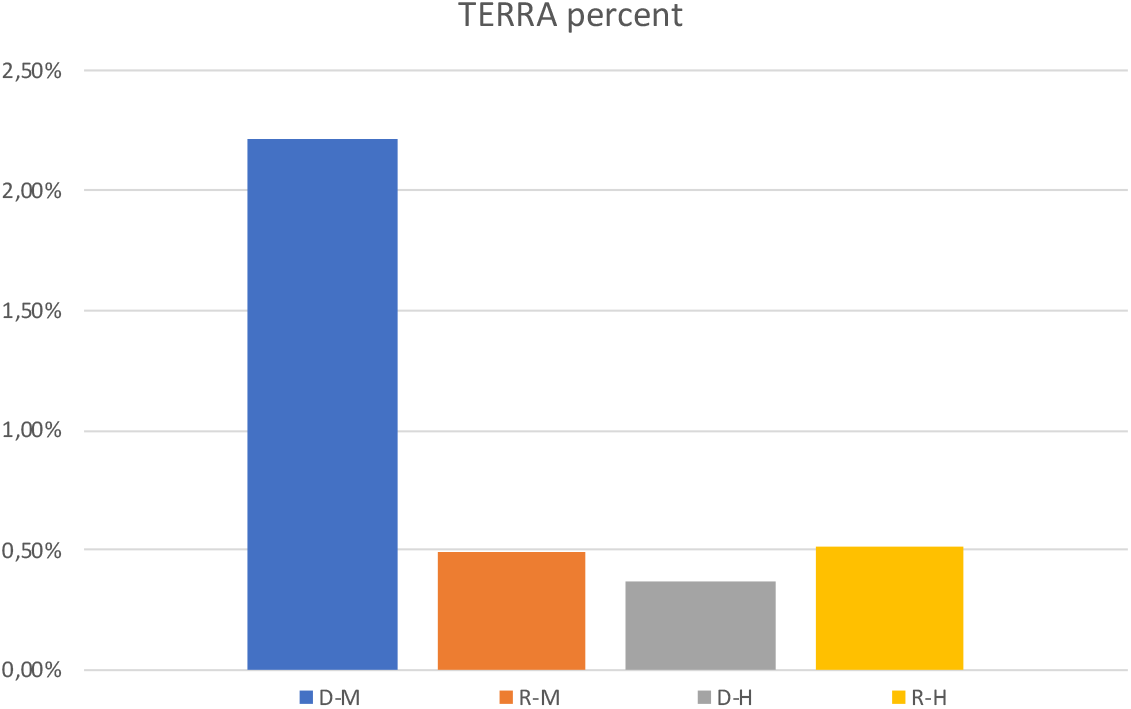
Part of Bioinformatics method for Supplementary Fig. 3 description: Reads with a length of less than 20 bp were discarded. Samples were then directly quantified using Salmon v0.12.0^32^ on an index built from the GRCm38 genome using GENCODE vM18. RNA composition estimation (Supplementary Fig. 3) were created by summing the reads of transcripts from the same RNA biotype and dividing by the total number of assigned reads in each sample^33^. **a**. RNA composition revealed by high-throughput RNA sequencing in sperm. Sperm transcriptome of free-RNA and DNA-bound RNA, resulting from individual mice (6 and 14 months old) reveals variable levels of transcripts mapping to protein-coding RNA, microRNA (miRNA), miscRNA, long intergenic non-coding RNA (lincRNA), sn/snoRNA and processed-transcript*. Y-axis shows the proportion of assigned reads in biological replicates of each sample group. **b**. RNA composition revealed by high-throughput RNA sequencing in testis. Transcriptome of testis of DNA-bound RNA (TD), resulting from individual mice (6 and 14 months old) reveals levels of transcripts mapping to protein-coding RNA, microRNA (miRNA), miscRNA, long intergenic non-coding RNA (lincRNA), sn/snoRNA and processed-transcript*. Y-axis shows the proportion of assigned reads in biological replicates of each sample group. **c**. Percent of TERRA* (minimum four consecutive CCCTAA repeats) revealed by high-throughput RNA sequencing of sperm of mouse DNA-bound RNA (D-M) and sperm human DNA-bound RNA (D-H) and free-RNA (R-H), Y-axis shows the TERRA percent of assigned reads.

**Supplementary Fig. 4.**
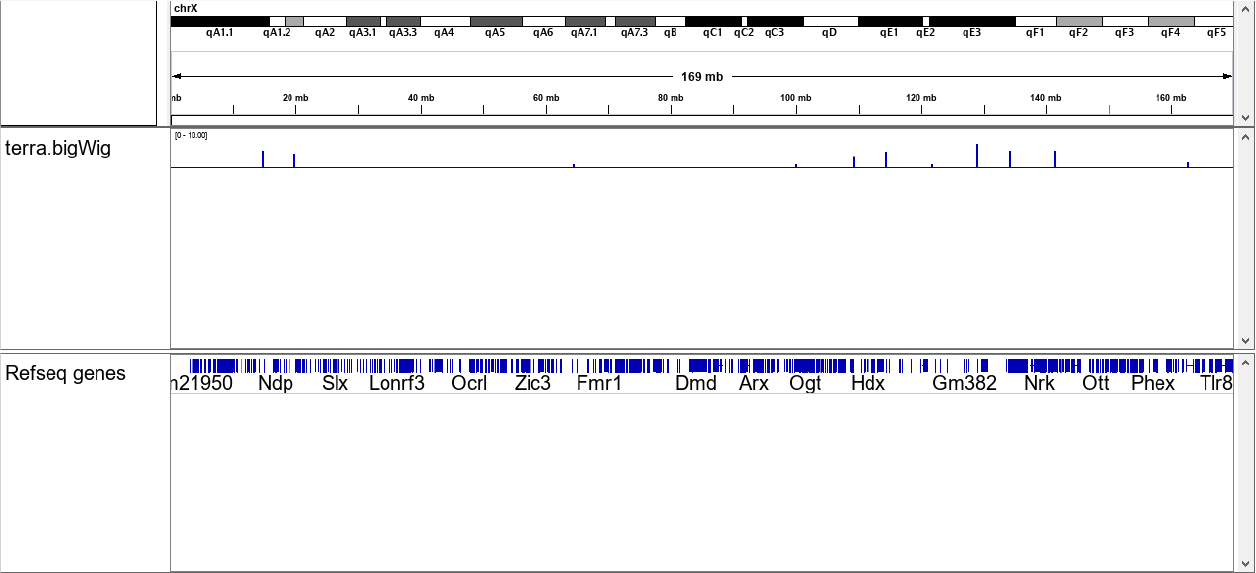
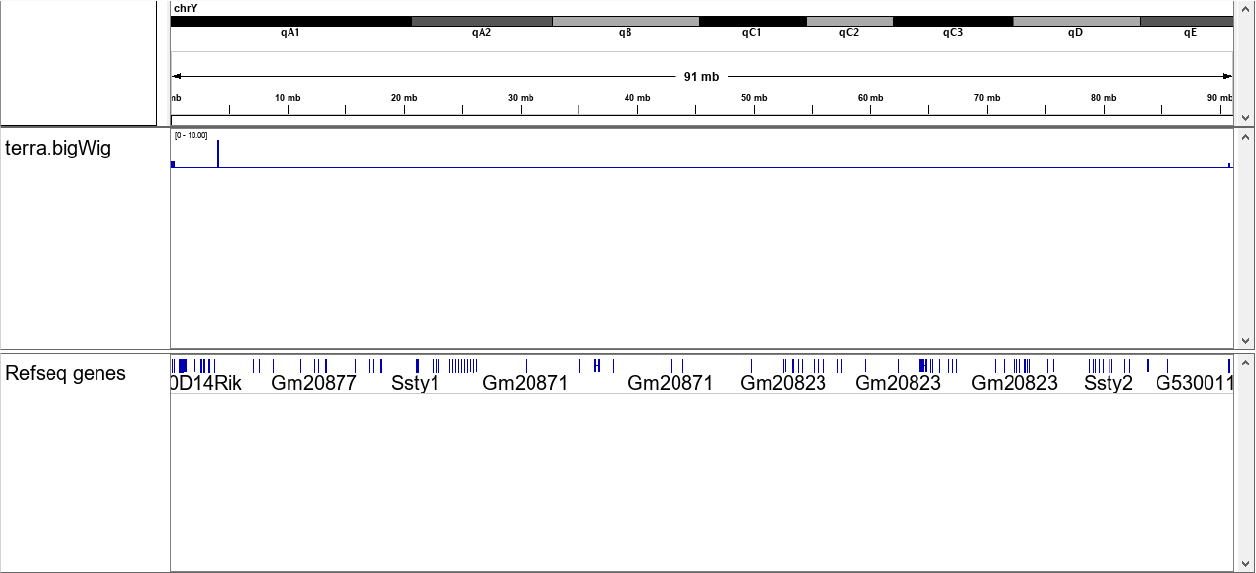
TERRA RNA spots are present in sex chromosomes (Fig 4a and b). Number of TERRA peaks in ChrX (Fig. 4a) is more than ChrY (Fig. 4b), however, the number of TERRA RNA foci are colocalized with pseudoautosomal regions. For instance, Asmt and Erdr1 according to Lee et al. Significant TERRA prominent peaks over ChrY in mouse sperm cells are associated with the ends of chromosome.

